# Overcoming efflux of fluorescent probes for actin imaging in living cells

**DOI:** 10.1101/2020.02.17.951525

**Authors:** Rūta Gerasimaitė, Jan Seikowski, Jens Schimpfhauser, Georgij Kostiuk, Tanja Gilat, Elisa D’Este, Sebastian Schnorrenberg, Gražvydas Lukinavičius

## Abstract

Actin cytoskeleton is crucial for endocytosis, intracellular trafficking, cell shape maintenance and a wide range of other cellular functions. Recently introduced cell-permeable fluorescent actin probes suffer from poor membrane permeability and stain some cell populations inhomogeneously due to the active efflux by the plasma membrane pumps. We addressed this issue by constructing a series of probes which employ modified rhodamine fluorophores. We found that the best performing probes are based on 6-carboxy-carbopyronine scaffold. These probes show preferential binding to F-actin, do not require efflux pumps inhibitors for staining and can be used for 2D and 3D fluorescence nanoscopy at high nanomolar concentrations without significant cytotoxicity. We demonstrate their excellent performance in multiple organisms and cell types: human cell lines, frog erythrocytes, fruit fly tissues and primary neurons.

## Introduction

Exploiting full potential of fluorescent microscopy requires tools and methods for specific labelling of targets in living cells.^1^ Small molecule probes, directed at organelles or proteins, are gaining momentum, as they do not require genetic modification and allow choice of fluorophore with desired optical properties.^2–5^ Such probes are synthesized by coupling a fluorophore to a well characterized, cell permeable ligand, which binds its target with high affinity and specificity. SiR-actin, consisting of 6’-carboxy silicon rhodamine (6’-SiR) **9** linked to the toxin jasplakinolide (JAS), is a widely used probe for staining actin in living cells.^6, 7^ Due to fluorogenicity, it yields high contrast confocal and stimulated emission depletion (STED) microscopy images. However, SiR-actin is susceptible to the active efflux by the plasma membrane pumps, which results in poor or mosaic staining of some cell lines.^6, 8, 9^ Herein we introduce new JAS-based probes for actin that are considerably less sensitive to the efflux and thus allow uniform staining of cancer cell lines that possess high multidrug resistance pump activity. In addition, we demonstrate that the fluorescent jasplakinolide derivatives are able to interact with both G-and F-actin.

The modular probes for live-cell imaging often exploit rhodamines as reporters.^10^ These dyes exist in a dynamic equilibrium between fluorescent zwitterionic state and non-fluorescent spirolactone. A more hydrophobic spirolactone facilitates passage through the cell membrane, whereas binding to the target shifts the equilibrium towards fluorescent form, thus ensuring high contrast images.^11^ Given this simple mechanism, most of the efforts to improve performance of modular probes were directed at fine-tuning spirolactone-zwitterion equilibrium by modifying the dye structure.^12, 13^ However, the optimized probes still required the use of verapamil to inhibit efflux pumps and to achieve efficient staining.^12^ This suggests that either the improvements were insufficient or other factors than spirolactone-zwitterion equilibrium might affect probe performance.

Previously, we and others have shown that finding an optimal fluorophore-ligand pair can make all the difference between useless and excellent-performing probe.^6, 14–16^ The changes as subtle as the attachment position on the dye can have a dramatic effect on staining of the target.^8, 16–18^ These examples call for a better understanding of interplay between the dye and the targeting ligand in determining the probe properties and underscore the importance to combinatorial screening when seeking the new probes. Thus, we explored the possibility to identify an improved living cell compatible actin probe.

## Experimental methods

### Spectra and fluorogenicity of the probes

Absorbance and fluorescence spectra of probes were recorded in four conditions: i) in phosphate buffered saline pH 7.4 (PBS), ii) in PBS with bovine serum albumin (BSA), iii) in General Actin Buffer (5 mM Tris-HCl pH 8.0, 0.2 mM CaCl_2_) with F-actin and iv) in PBS with 0.1% SDS. The following reagents were mixed in total volume of 250 μl in a black glass-bottom 96-well plate (MatTek, Cat. No. PBK96G-1.5-5-F): respective buffer, 2 μM probe, 0.5 mg/ml actin or BSA. Then, 25 μl of 10× actin polymerization buffer (actin sample) or PBS (all other samples) was added, the solution was mixed by pipetting and the plates were incubated for 1 h at 37°C. (10×actin polymerization buffer contains 500 mM KCl, 20 mM MgCl2, 0.05 mM guanidine carbonate and 10 mM ATP). The spectra were acquired using Tecan Spark20M plate reader in a bottom reading mode. The spectra were processed and analyzed with a|e v2.2 (background subtraction and normalization) (FluorTools, http://www.fluortools.com) and Spectragryph v1.2.11 (spectra averaging) (F.Menges “Spectragryph - optical spectroscopy software”, Version 1.2.11, 2019, http://www.effemm2.de/spectragryph/). To account for light scattering, spectra of the solutions containing no probes, but equivalent amount of DMSO were acquired and subtracted from the respective probe spectra. Then the absorbance and fluorescence spectra of a probe was normalized to the A_max_ or F_max_ of the SDS sample. The experiment was repeated three times, the normalized spectra were averaged and are presented in Fig. S1.

### Maintenance of cells

U-2 OS and Human primary dermal fibroblasts and HeLa cells were cultured in high-glucose DMEM (Dulbecco’s Modified Eagle Medium, Life Technologies, Cat. No. 31053-028) supplemented with GlutaMAX-1 (Life Technologies, Cat. No. 35050-038) and 10% foetal bovine serum (FBS, Life Technologies, Cat. No. 10270-106) in a humidified 5% CO2 incubator at 37 °C.

Normal neonatal human melanocytes (MatTek Corporation, Cat. NHM-CRY-NEO) were cultured in Normal human melanocyte cells growth medium (MatTek Corporation, Cat. NHM-GM) in a humidified 5% CO2 incubator at 37 °C.

All cell types were split every 3-4 days or at confluence. Staining performed in the growth media.

### Staining and imaging of living cells

U-2 OS cells were grown in 12-well uncoated glass bottom plates (MatTek) in DMEM supplemented with 10 % FBS. For staining, the cells were incubated for 2 h at 37°C in DMEM containing 250 nM or 1 μM probe and 0.1 μg/ml Hoechst in the presence or absence of 10 μM verapamil. After the incubation, the cells were washed once with DMEM and once with Hank’s Balanced Salt Solution (HBSS). Fresh DMEM was added and the cells were imaged on a wide-field Lionheart FX Automated Microscope (Biotek) with 20× objective, using laser autofocus. 16 fields of view in 3 focusing planes, spanning 6 μm in thickness, were acquired per well. Image stitching and focus stacking was performed using in-built Gene 5 software (Biotek). The final images encompassed 1380×950 μm field of view and contained at least 220 and up to 2000 cells, which ensured analysis with no bias towards a well-stained sub-population. Each experiment was repeated independently 3 times. Actin staining was quantified with automated pipeline built in CellProfiler v.3.1.8 (Fig. S2).^19^

### Formaldehyde fixation and determination of *K_D_^F-actin^*

U-2 OS cells were grown as before, briefly washed with 1 ml HBSS and incubated for 10 min. with 1 ml of 4% formaldehyde in PBS (pH 7.4). The solution was removed, and 1 ml of 30 mM glycine in PBS (pH 7.4) was added for 5 min. Then the samples were washed 3× 5 min. with 1 ml PBS, incubated for 5 min. with 1 ml 0.1% reduced Triton X-100 in PBS and washed again 3× 5 min. with 1 ml PBS. The samples were blocked by incubating with 1 ml 1% BSA in PBS for 30 min. and stored in PBS at 4°C. The cells were stained by incubating with 1 ml of 1-1000 nM probe and 0.1 μg/ml Hoechst in PBS for 30 min. at RT, washed 2 times with 1 ml PBS and imaged on a wide-field Lionheart FX Automated Microscope (Biotek) with 4× objective, using laser autofocus. Actin staining was measured using CellProfiler v.3.1.8 (ref. ^19^) pipeline, as explained in Fig. S2, and the data was fitted into a dose response equation (1):

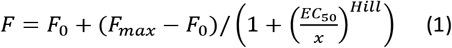

where *F_o_* – signal without probe, *F_max_* – maximum staining, x – probe concentration, *Hill* - Hill slope coefficient determining the steepness of a dose-response curve, *EC_50_* - probe concentration that results in half-maximum staining (*F_max_-F_0_*). As Hill coefficient was found to be close to 1, and probe concentration is >> actin concentration, *EC_50_* is equivalent to apparent *K_D_^F-actin^*.

### Probe interaction with G-actin

The reactions were performed in General Actin Buffer, supplemented with 0.2 mM ATP and 0.5 mM DTT. Eleven 2-fold dilutions of actin were prepared, starting from 8 μM, and aliquots of 25 μl were transferred into a black 96-well half-area plate (Greiner Bio-One, Cat. No. 675076). Immediately before use, probes were diluted from DMSO stock solution to 20 nM in the same buffer at room temperature, and 25 μl aliquots were mixed with actin dilution. The samples were incubated at 37°C for 1 h and the fluorescence was read with Tecan Spark20M plate reader in a top-reading mode. Excitation/emission wavelengths (nm) were as follows: 470/530 for LIVE 510 and LIVE 515, 510/560 for 530RH, 570/610 for 580CP, 570/630 for 610CP and 590/650 for SiR; excitation and emission bandwidths were in all the cases 15 nm. Background fluorescence of the sample without actin was subtracted. When assessing effects of latrunculin A or DNase I, actin dilutions were prepared in the buffer containing 10 μM of these inhibitors, thus making their concentration constant in the whole titration series and equal to 5 μM in the final reaction mixture. The data was fitted into a full equation (2) of single site binding:

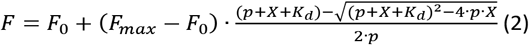

where *F*_0_ – fluorescence of probe without target, *F*_max_ – fluorescence of probe at saturating concentration, *p* – probe concentration, X – actin concentration, *K_d_* – dissociation constant of the probe. All measurements were performed 3-5 times on different days.

### STED nanoscopy of neurons

Primary rat hippocampal neurons were prepared as previously described.^6^ Cells were labelled in growth medium at 30 days *in vitro* for 45 min. with 100 nM probe and washed prior to imaging. Axons were identified by labelling with anti-neurofascin antibody (NeuroMab, 75-172) and an anti-mouse AlexaFluor 488 secondary antibody.^6^ Imaging was performed in ACSF buffer on an Abberior easy3D STED/RESOLFT QUAD scanning microscope (Abberior Instruments GmbH, Göttingen, Germany) built on a motorized inverted microscope IX83 (Olympus, Tokyo, Japan). The microscope is equipped with pulsed STED lasers at 595 nm and 775 nm, and with 355 nm, 405 nm, 485 nm, 561 nm, and 640 nm excitation lasers. Spectral detection was performed with avalanche photodiodes (APD) in the following spectral windows: 650-800 nm and 505-550 nm. Images were acquired with a 100x/1.40 UPlanSApo Oilimmersion objective lens (Olympus) and pixel size was 30 nm.

### Confocal, 2D and 3D STED imaging

Confocal imaging was performed on a Leica SP8 (Leica Microsystems, Mannheim, Germany) inverted confocal microscope equipped with an HC PL APO CS2 63x/1.40 OIL objective. Images were acquired using a 1000 Hz bidirectional scanner, a voxel size of 80 nm × 80 nm x 1000 nm, a pinhole of 1 AU and line averaging of 3. Hoechst 33342 was excited with a 405 nm laser and detected with a regular PMT in the 430–480 nm range. Fluorescent actin probes were excited and detected using the following parameters: LIVE 515 probe – excited with 488 nm laser and detected with Leica HyD detector set within the spectral range of 530–580 nm, 580CP probes – excited with 561nm laser and detected in the range 610–660 nm, 610CP or Sir probes – exited with 633 nm laser and detected in the range 650–700nm.

Comparative confocal and STED images were acquired on Abberior STED 775 QUAD scanning microscope (Abberior Instruments GmbH, Germany) equipped with 561 nm and 640 nm 40 MHz pulsed excitation lasers, a pulsed 775 nm 40 MHz STED laser, and an UPlanSApo 100x/1.40 Oil objective. The following detection windows were used: 580CP channel 615 / 20 nm and 610CP/SiR channel 685 / 70 nm. In this setup, pixel size was 30 nm in xy plane, pinhole was set to 1 AU for 2D STED images. The laser power was optimized for each sample. 3D STED images were acquired using pinhole set to 0.8 AU, voxel size set to 40 x 40 x 40 nm, 3D STED doughnut set to 90%, single line accumulation and xzy scanning mode. Acquired images were processed using Fiji^20^ and SVI Huygens deconvolution software.

### Airyscan microscopy

Freshly extracted frog erythrocytes were diluted 1000× in RBC buffer (10 mM HEPES pH 7.4, 150 mM NaCl, 0.1% glycerol) supplemented with 250 nM 6-610CP-JAS, 200 μl aliquots were seeded into 10-well plate (Greiner bio-one culture slides, PS, 75/25 mm. (Art. Nr.: 543079)) and incubated for 2 h at room temperature. Imaging was performed on Zeiss LSM880 system equipped with oil immersion Plan-Apochromat 63X/1.40 Oil Corr M27 objective (Carl Zeiss) at room temperature. Stained erythrocytes were imaged in Z-stack mode according Nyquist parameters (x × y × z = 55 × 55 × 150 nm). 633 nm diode laser was used for excitation, emission was detected using BP 570 - 620 + LP 645 filter onto Airyscan fast detector. The signal was averaged from 2 frame scans. Live-cell imaging was performed on the same system equipped with incubator set to 37°C and 5% CO_2_. The images were reconstructed using Zen software (Carl Zeiss) and processed with Fiji^20^.

Human fibroblasts were stained in a 10-well plate (Greiner bio-one culture slides, PS, 75/25 mm. (Art. Nr.:543079) with 200 μl of DMEM supplemented with 10% FBS and 250 nM 6-610CP-JAS for 2h at 37°C and 5% CO2. Imaging was performed on Zeiss LSM880 system equipped with oil immersion Plan-Apochromat 63X/1.40 Oil Corr M27 objective (Carl Zeiss) at 37°C and 5% CO2. Stained primary human fibroblasts were imaged in time-lapse mode for 15 min. every 15 s in one Z-plane (x × y = 55 × 55 nm). 633 nm diode laser was used for excitation, emission was detected using BP 570 - 620 + LP 645 filter onto Airyscan fast detector. The signal was averaged from 4 frame scans. The images were processed with Fiji^20^.

### Maintenance and preparation of Drosophila melanogaster larvae

Wildtype OregonR Drosophila melanogaster were raised on standard cornmeal-yeast-agar-medium at 25 °C and used for all experiments. For staining of living Drosophila melanogaster tissues, wandering third instar larvae were dissected in 1x PBS pH 7.4 (Phosphate Buffered Saline) (Life Technologies, Carlsbad, California USA) and the inverted front half of the larva was incubated with probes of 1 μM concentration in 1x PBS for 1 h at room temperature. After single washing step with 1x PBS, isolated tissues were mounted in Schneider Drosophila Medium (ThermoFischer Scientific) under a coverslip and sealed with nontoxic duplicating silicone (picodent, Wipperfuerth, Germany) to prevent evaporation of the medium during imaging.

## Results and discussion

### Design and synthesis of actin probes

We prepared jasplakinolide *N^ω^*-Boc-lysine analogue ^7, 21^ (JAS-Boc **1**) and conjugated it to a series of STED nanoscopy compatible fluorophores – 6’-530RH, 5’-580CP, 5’-610CP, 6’610CP and compared them to SiR-actin (Fig. 1a). In order to gain a better insight into the structure-performance relationship, we also analyzed recently published 6-LIVE 510-JAS and 6-LIVE 515-JAS ^22^ and 6’-580CP-JAS ^23^ probes.

**Fig. 1.**
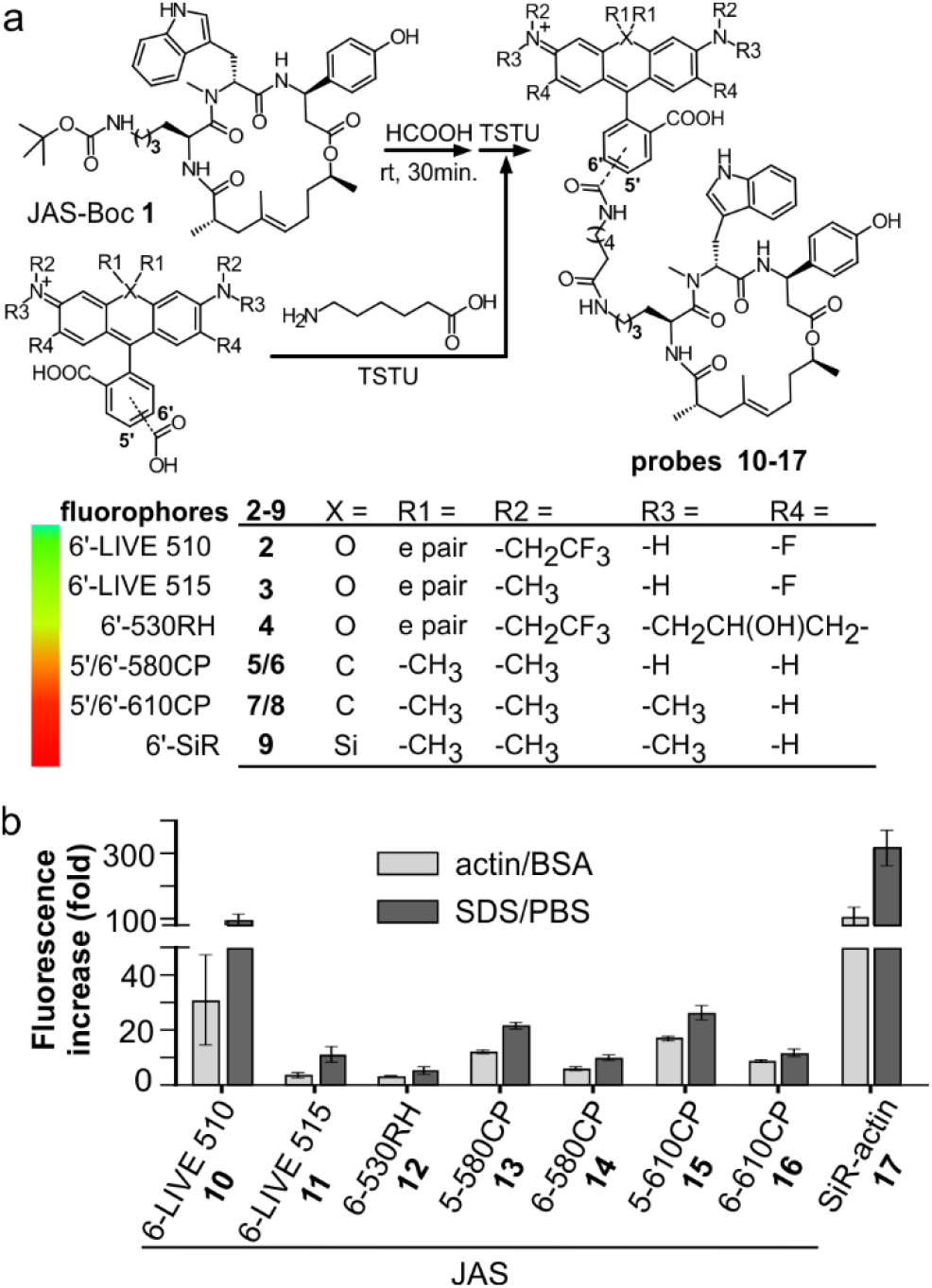
Structure and optical properties of the new actin probes. (a) Simplified synthesis scheme and chemical structures of fluorophores and actin probes characterized in this work. (b) Fluorescence increase of 1.6 μM probes upon incubation with 10 μM F-actin or 0.1% SDS. Mean ± SD, N=3.

### Chromogenicity and fluorogenicity of the probes

All probes were chromogenic and fluorogenic, i.e. showed absorbance and fluorescence increase upon SDS detergent addition and actin binding (Fig. 1b and Fig. S1a-h). Probe’s optical properties in actin-bound versus free state determine the image contrast and possibility of no-wash imaging. SDS dissolves the aggregates, shifts equilibrium to the fluorescent zwitterionic state and thus allows measuring maximum fluorescence and absorbance values in aqueous environment.^11^ BSA was used to estimate fluorescence increase due to unspecific interactions, which was negligible in all the cases (Fig. S1). Fluorogenicity in the presence of actin or SDS followed the same trend: SiR-actin and 6-LIVE 510-JAS were the most fluorogenic, while 6-530RH-JAS and 6-LIVE 515-JAS were the least fluorogenic (Fig. 1b). In agreement with the previous studies, high fluorogenicity was determined by a low fluorescence in the unbound state, rather than fluorescence enhancement after interaction with the target (Fig. S1).

Most probes achieved ~ 80% of the maximal absorbance and fluorescence, when bound to actin (Fig. S1i). Exceptionally, 6-LIVE 510-JAS and 6-LIVE 515-JAS gained ~80% in absorbance, indicating that the binding had switched the equilibrium towards zwitterionic state, but fluorescence increase reached only 30-50% of that in SDS. This suggest fluorescence quenching due to the interaction with actin or aromatic residues of jasplakinolide.^24^

### Staining actin in living cells

U-2 OS cells display drug efflux activity, thus their uniform and efficient staining with rhodamine probes requires verapamil.^8^ We used these “difficult” cells to reveal performance differences among the new probes by quantifying the actin staining in the absence and in the presence of verapamil (Fig. S3). The ratio of the measured values close to 1 indicates that the active efflux does not limit actin staining. Indeed, the probes behaved very differently (Fig. 2a and Fig. S3a). As previously reported, SiR-actin, stained the cells brightly in the presence of verapamil ^6^, but only a sub-population of cells was stained without verapamil. 6-LIVE 510-JAS, 5-580CP-JAS and 5-610CP-JAS performed similarly. 6-530RH-JAS stained cells weakly, required higher concentration and produced numerous fluorescent aggregates. Surprisingly, 6-LIVE 515-JAS, 6-580CP-JAS and 6-610CP-JAS stained the cell population uniformly and verapamil addition only moderately increased the staining intensity (Fig. 2a, b). Thus, jasplakinolide-based actin probes represent yet another example where isomerism of fluorophores is decisive to probe performance. We found no correlation between high fluorogenicity and good performance in the present series of probes. In fact, our best performing probes, 6-580CP-JAS, 6-610CP-JAS and 6-LIVE 515-JAS, are among the least fluorogenic (Fig. 1b). There was no difference between staining by 5’-and 6’-isomers of 580CP-JAS and 610CP-JAS in the presence of verapamil (Fig. 2c). Thus improved staining by the 6’-isomers might be caused by their impaired efflux.

**Fig. 2.**
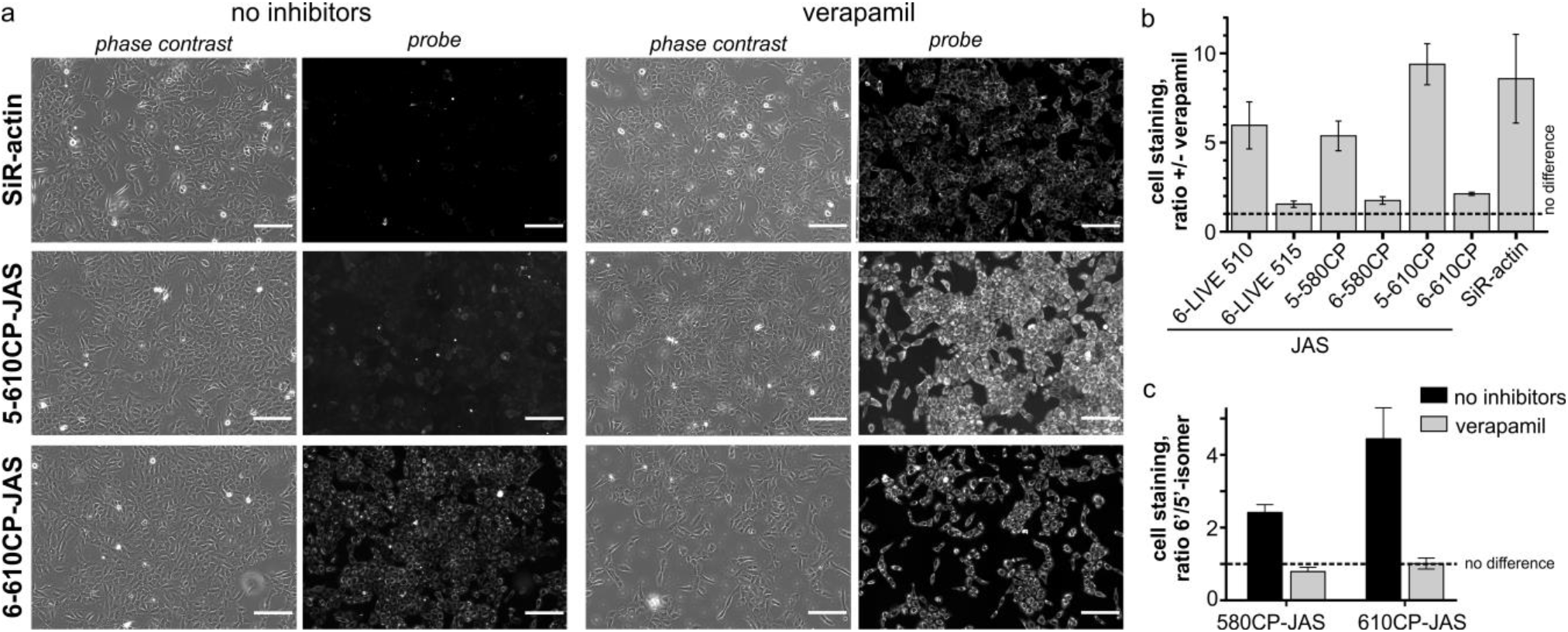
Requirement of verapamil for efficient staining of actin in U-2 OS cells. (a) Living cells were incubated with 250 nM SiR-actin, 5-610CP-JAS or 6-610CP-JAS for 2 h at 37°C in the presence or absence of 10 μM verapamil, washed and imaged on the automated wide-field microscope. Scale bar – 200 μm. (b) Enhancement of staining by verapamil. (c) A difference between 5’-and 6’-isomers of the probes is observed only in the absence of verapamil. (b), (c) Actin staining was quantified with Cell Profiler pipeline, as described in (Fig. S2). N = 3, at least 200 cells per conditions were measured in each of three independent experiments. Data is presented as mean ± SD.

### Interaction with F-actin

To asses binding affinities, we stained formaldehyde-fixed and permeabilized U-2 OS cells with 1 - 1000 nM probes and quantified the resulting fluorescence in the cytoplasm. The data fitted well into a dose response equation with Hill coefficient equal to 1, thus the derived *EC_50_^staining^* is equivalent to apparent *K_D_^F-actin^* (Fig. S4 and Table 1). Interestingly, *K_D_^F-actin^* of all probes was very similar and fell in the range of 15 - 60 nM, which is very close to the reported *K_D_^F-actin^* = 15 nM of jasplakinolide.^25^

**Table 1.**
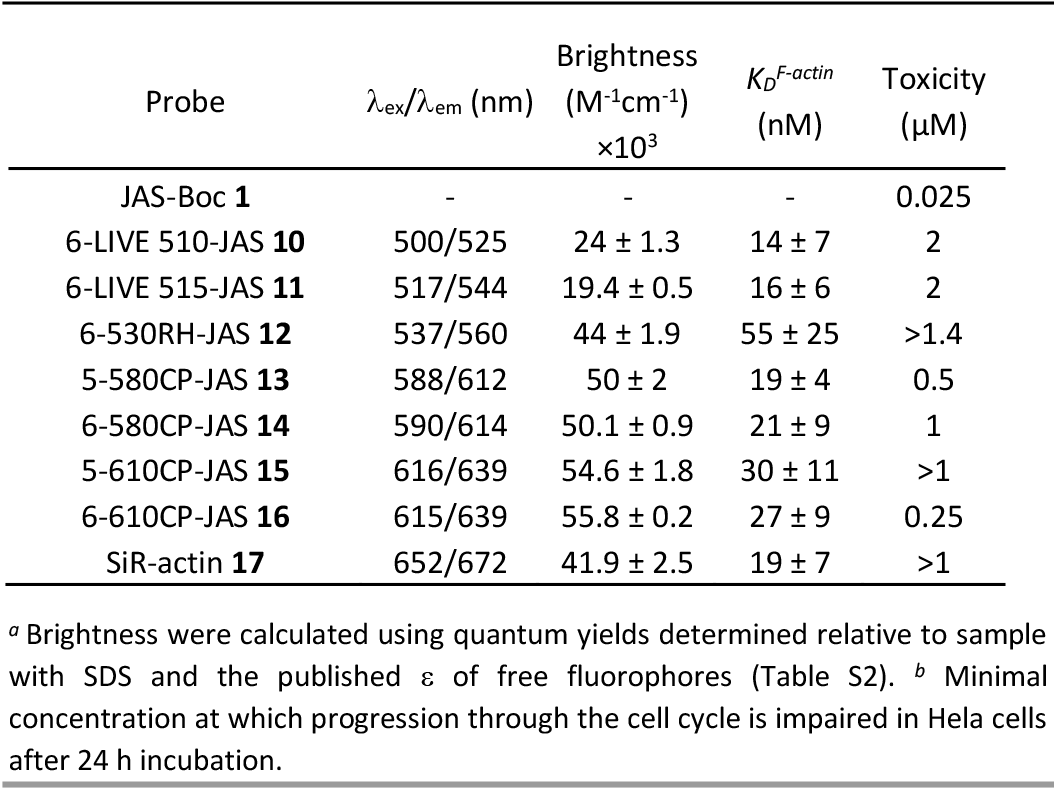
Properties of the fluorescent actin probes.

This indicates that attachment of fluorophore does not change significantly jasplakinolide affinity to actin. Thus differences in probe performance cannot be explained by differences of interaction with actin, but arise from differences in the cell entry and/or retention.

### Cytotoxicity of the probes

While SiR-actin is not toxic over a wide concentration range^6^, the parent compound, jasplakinolide, is a powerful toxin.^26^ We assessed cytotoxicity of the new probes by measuring cell cycle perturbations in Hela cells after 24 h incubation (Fig. S5): JAS-Boc showed toxicity threshold at 25 nM and for all new probes it was at ~1 μM (Table 1). The better staining probes were slightly more toxic, which might reflect their reduced efflux. However, even the most toxic probes (250 nM) were ~10-fold less potent compared to the parent compound. This indicates that cell entry remains a limiting step in staining actin. Rhodamines are known substrates for P-glycoprotein 1 - one of the major players in ATP-dependent efflux.^27^ Limited passive permeability and active efflux can compromise staining and toxicity by reducing intracellular probe concentra-tion.^28^ This is well illustrated by extreme dependency of SiR-actin staining on verapamil.

### Interaction with G-actin

Jasplakinolide binds and stabilizes F-actin filaments.^25^ Fluorogenic nature of the probes allowed us to investigate their interaction with G-actin. We saw a clear increase in fluorescence while titrating 5-610CP-JAS, 6-610CP-JAS and SiR-actin with G-actin in a low salt buffer that does not support polymerization (Fig. S6). The apparent *K_D_* values for 5/6-610CP-JAS were surprisingly close to *K_D_^F-actin^* (Fig. 3a). To ensure that in our assay actin remains unpolymerized, we repeated titrations in the presence of polymerization inhibitors –toxin latrunculin A (LatA)^29^ and DNase I which bind to the sites non-overlapping with jasplakinolide binding site.^25, 30, 31^ LatA had no effect on *K_D_*, but in the presence of DNase I *K_D_* of all three probes increased by ~10-fold (Fig. 3a).

**Fig. 3.**
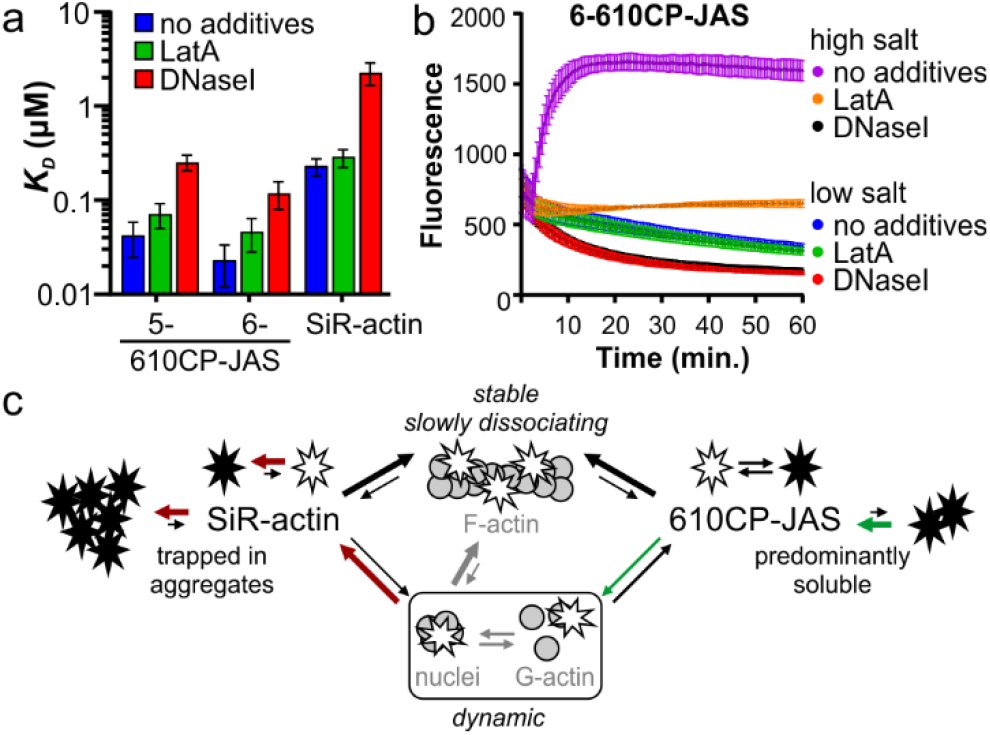
Interaction of probes with G-actin. (a) *K_D_* values derived from titrating 10 nM probes with actin under low salt (non-polymerizing) conditions. Mean ± SD; N = 5 without additives, N = 3 in the presence of polymerization inhibitors. (b) Time-courses of polymerization of 4 μM pyrene actin in the presence of 10 nM 6-610CP-JAS without additives, with 5 μM latrunculin A or with 5 μM DNase I. N = 3; mean ± SD. (c) Model explaining higher selectivity of more hydrophobic probe to F-actin. Aggregation strongly affects lower affinity dynamic interaction with G-actin, but has little impact on more stable interaction with F-actin.

To resolve this discrepancy, we tested for possible actin polymerization by monitoring fluorescence of pyrene-labelled G-actin under the conditions of our titration (Fig. 3b). As the rate of nucleation depends on G-actin concentration^32^, we performed this experiment at the highest G-actin concentration (4 μM) used. The addition of high salt buffer to the samples induced rapid actin polymerization which was completely blocked by DNase I and largely inhibited by LatA. Slow fluorescence decrease was observed in a low salt buffer used for the titrations, which approached plateau after 1 h of incubation. Addition of LatA had no effect, while DNase I accelerated this process. Because of a low concentration (10 nM) probes had no effect on these processes. Taken together, this data indicates that a fraction of actin in a low salt buffer exists in a multimeric state that is completely depolymerized by DNaseI.^33^ Presence of highly polymerized filaments is unlikely, because actin stock was centrifuged immediately before the experiment, but G-actin can exist under the rapid equilibrium with nucleation centres and/or short filaments.

Therefore we assume that *K_D_* determined in the presence of DNase I represents the true affinity of the probes to G-actin (*K_D_^G-actin^*).

Comparison of *K_D_^G-actin^* (Fig. 3a) and *K_D_^F-actin^* (Table 1) reveals that 610CP-JAS has ~10-fold preference, while SiR-actin has ~100-fold preference for F-actin. We explain with difference by a higher aggregation propensity of SiR-actin.^8, 14^ Assuming that interaction of jasplakinolide probes with G-actin (and very short filaments) is more dynamic and has higher *k_off_* rate, it would be more sensitive to aggregation than a more stable interaction with F-actin. This effect is stronger for SiR-actin than for 610CP-JAS probes, resulting in a higher preference of SiR-actin to F-actin *in vitro* (Fig. 3c). However, in living cells the concentrations of both, G-and F-actin, is in high micromolar range^34, 35^, indicating that both forms can be stained by both probes. This is supported by the microscopy imaging showing highly dynamic diffuse cytoplasm staining in addition to the bright F-actin fibres (Video S1).

### Imaging actin in living cells and tissues

Our new probes are compatible with various nanoscopy setups and can be used to stain variety of samples with high specificity for actin (Fig. 4). 6-580CP-JAS can be combined with SiR-based probes for two colour STED imaging in cell cultures and in living tissues, where small probe size ensures efficient volume penetration. Efficient staining by 6-610CP-JAS allows imaging actin in erythrocytes that contain high haemoglobin absorption. Also, it was successfully used for 3D isotropic STED imaging of actin in human fibroblasts (Video S2).

**Fig. 4.**
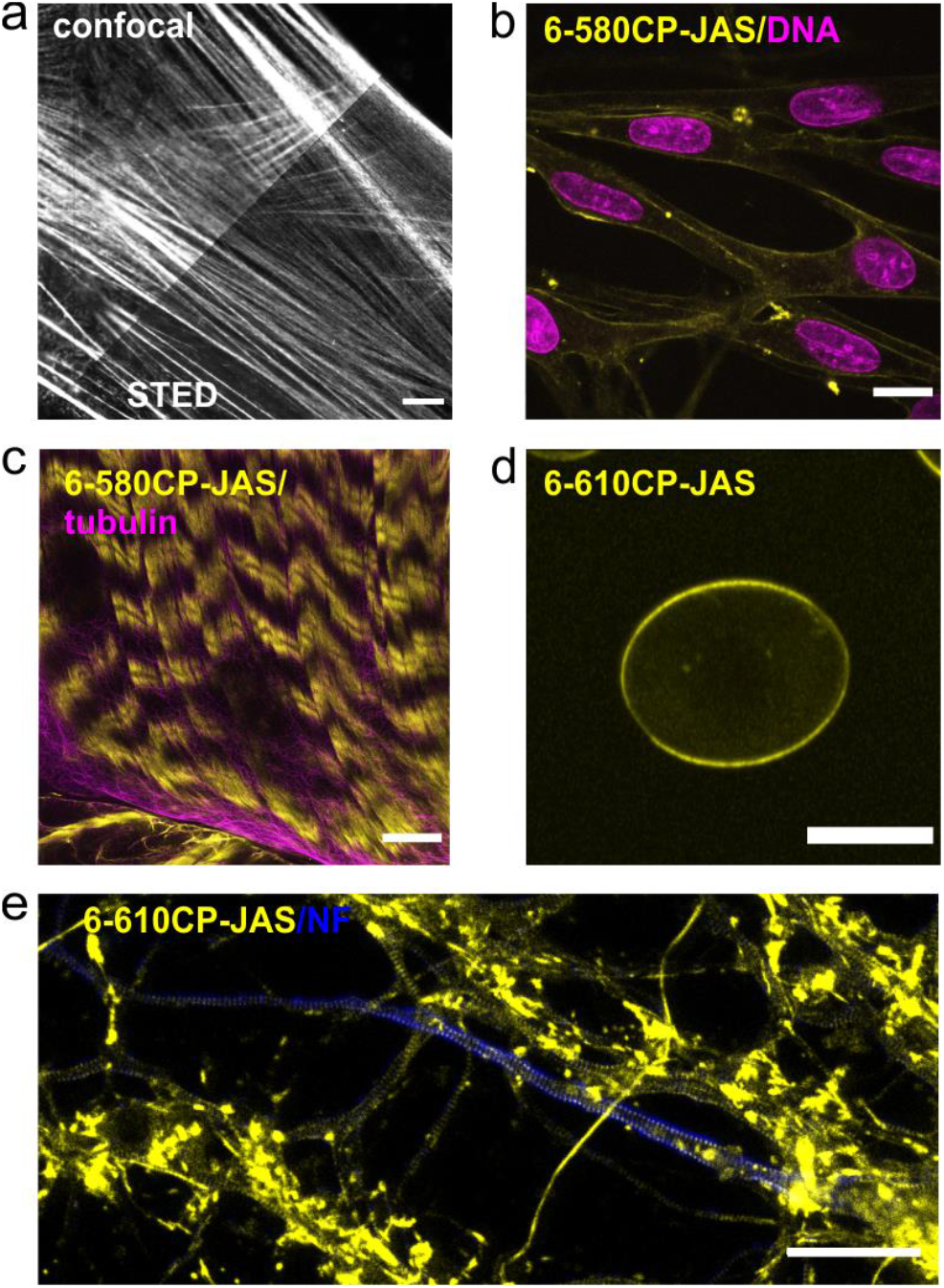
Confocal and nanoscopy images of living cells and tissues stained with the new actin probes. (a) Comparison of confocal and STED images of human fibroblasts stained with 6-610CP-JAS. (b) Human melanocytes co-stained with 6-580CP-JAS and 5-SiR-Hoechst.^8^ (c) live-cell imaging of body wall muscle of dissected *D. melanogaster* larva costained with 6-580CP-JAS and 6-SiR-CTX.^14^ (d) Max intensity projection of a frog erythrocyte stained with 6-610CP-JAS. (e) Rat primary neuron culture co-stained with 6-610CP-JAS and neurofascin (AlexaFluor 488). Scale bars 10 μm (a-d), 5 μm (e).

## Conclusions

In summary, we demonstrate that subtle differences in probe structure might affect their interaction with efflux pumps, which can dramatically affect probe performance in living cells. 6-LIVE 515-JAS, 6-580CP-JAS and 6-610CP-JAS represent a valuable addition to the actin imaging toolbox as they can be used for staining cells with high efflux pump activity without inhibitors. Notably, all probes are able to bind both G-and F-actin which should be taken into account while interpreting *in vitro* and *in vivo* observations. The membrane permeability remains a limiting factor and work directed at its improving is currently under way.

## Acknowledgements

The authors acknowledge funding of the Max Planck Society.

G. L. is grateful to the Max Planck Society for a Nobel Laureate Fellowship. G.K. acknowledges Max Planck Institute for Biophysical Chemistry for Manfred Eigen Fellowship. The authors thank dr. Vladimir Belov and dr. Jonas Bucevičius for the discussions and critical reading of the manuscript.

