## Supplementary material for "Overcoming efflux of fluorescent probes for actin imaging in living cells"

#### Supplementary Figures

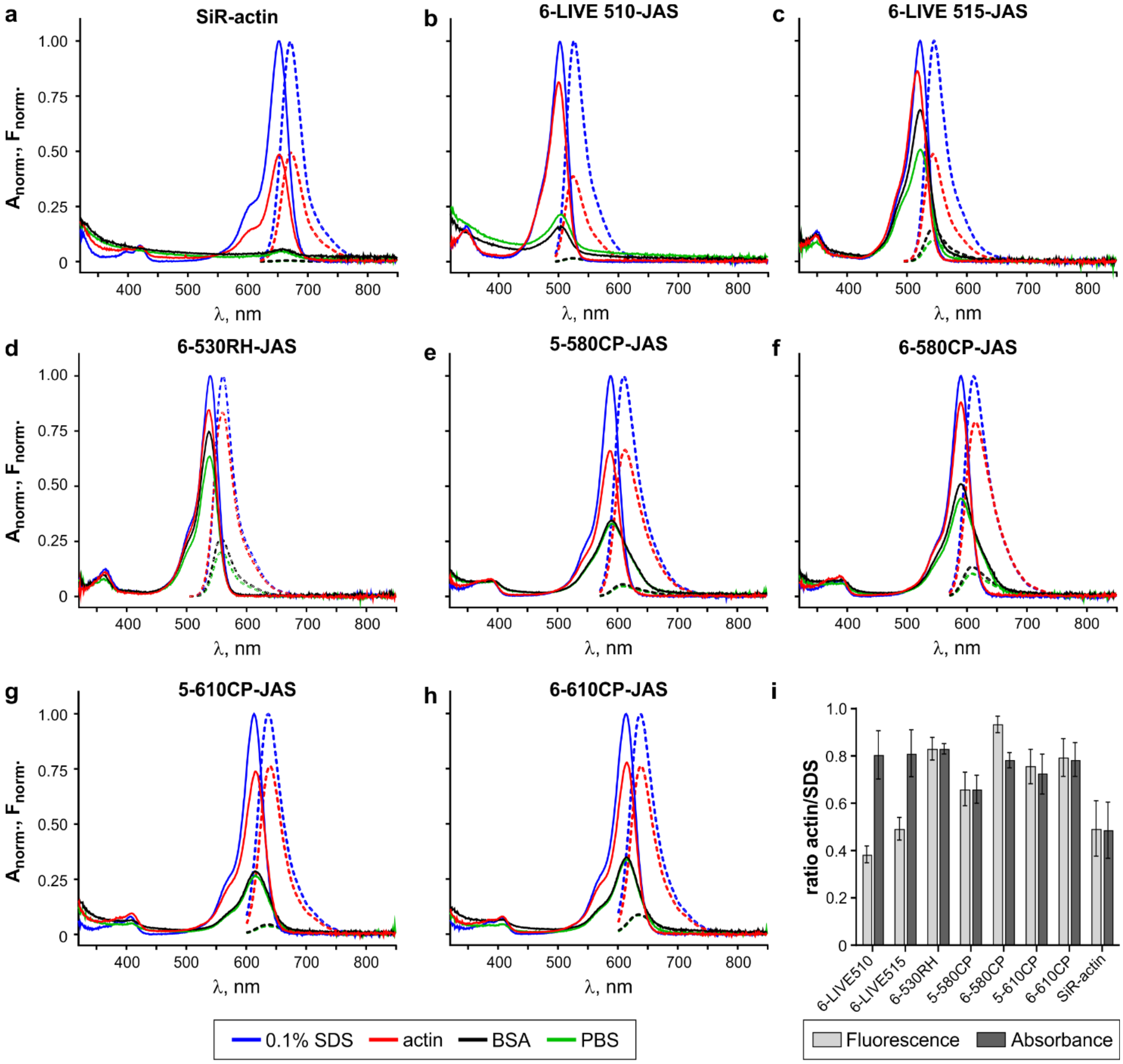

**Figure S1. Absorbance and fluorescence spectra of actin probes.**

**a-h**, normalized absorbance (solid line) and fluorescence (dotted line) spectra of the indicated probes. **i**, ratio of fluorescence or absorbance in the presence of 1.6 µM actin and respective values in the presence of 0.1% SDS. Spectra are presented as average of three independent measurements.

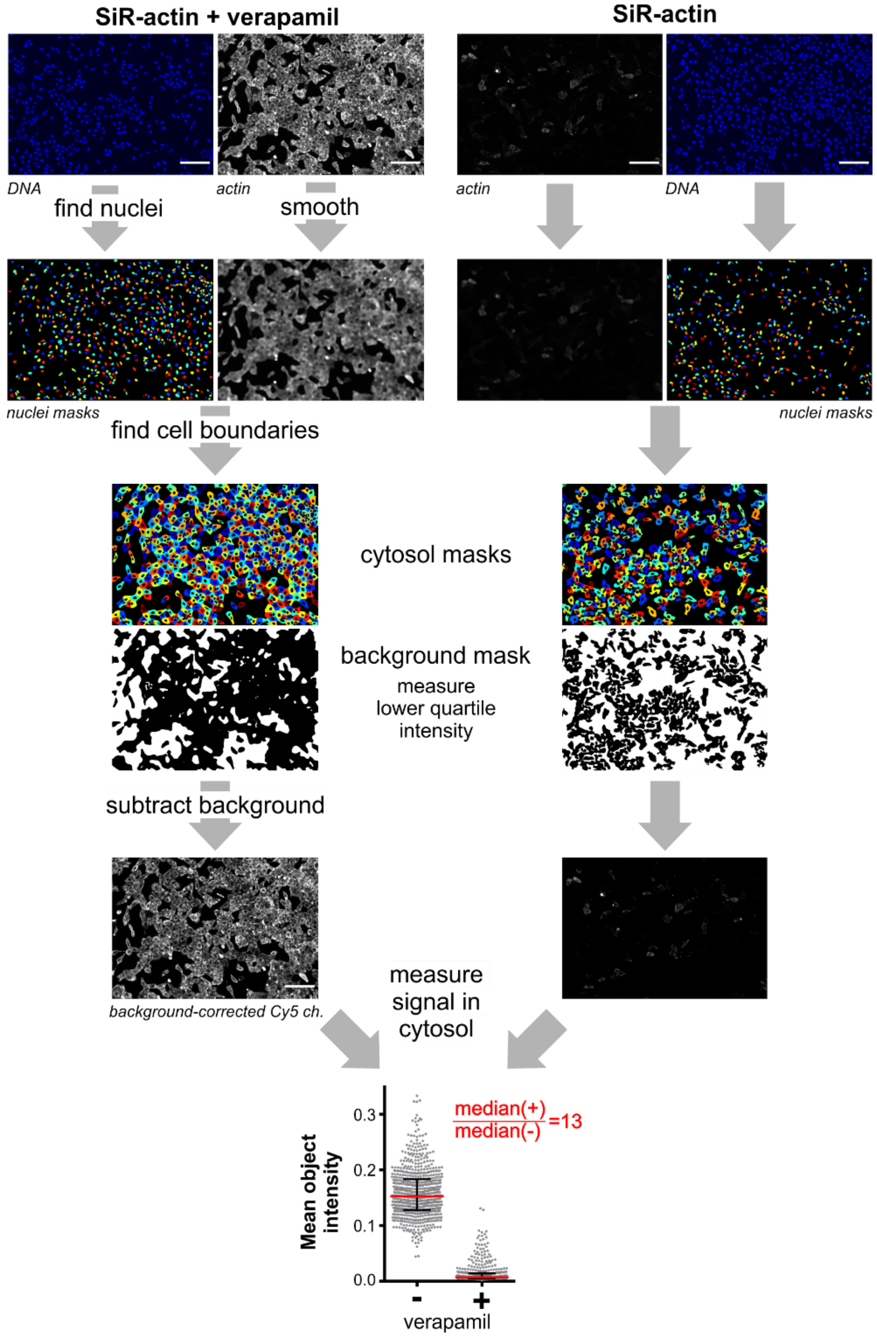

**Figure S2. Major steps in actin staining quantification pipeline.**
Example images of U2OS cells stained with 1 µM SiR-actin in the presence or absence 10 µM verapamil, scale bar 200 µm. Nuclei are identified in the Hoechst 33342 channel and used as seeds to detect cells in the smoothed probe channel. Cytosol mask is generated by subtracting nucleus from the cell. Background is defined as lower quartile intensity of the areas not covered by the cells. Background is subtracted from the original probe channel and mean pixel intensity is measured within the cytosol mask of each cell. 200-2000 cells are analyzed per image and median object pixel intensity of the image is taken as a readout of actin staining. Ratio of staining in the presence versus in the absence of verapamil is taken as a measure of the probe's susceptibility to active efflux. The derived results are presented in main text Figure 2(b) and (c).

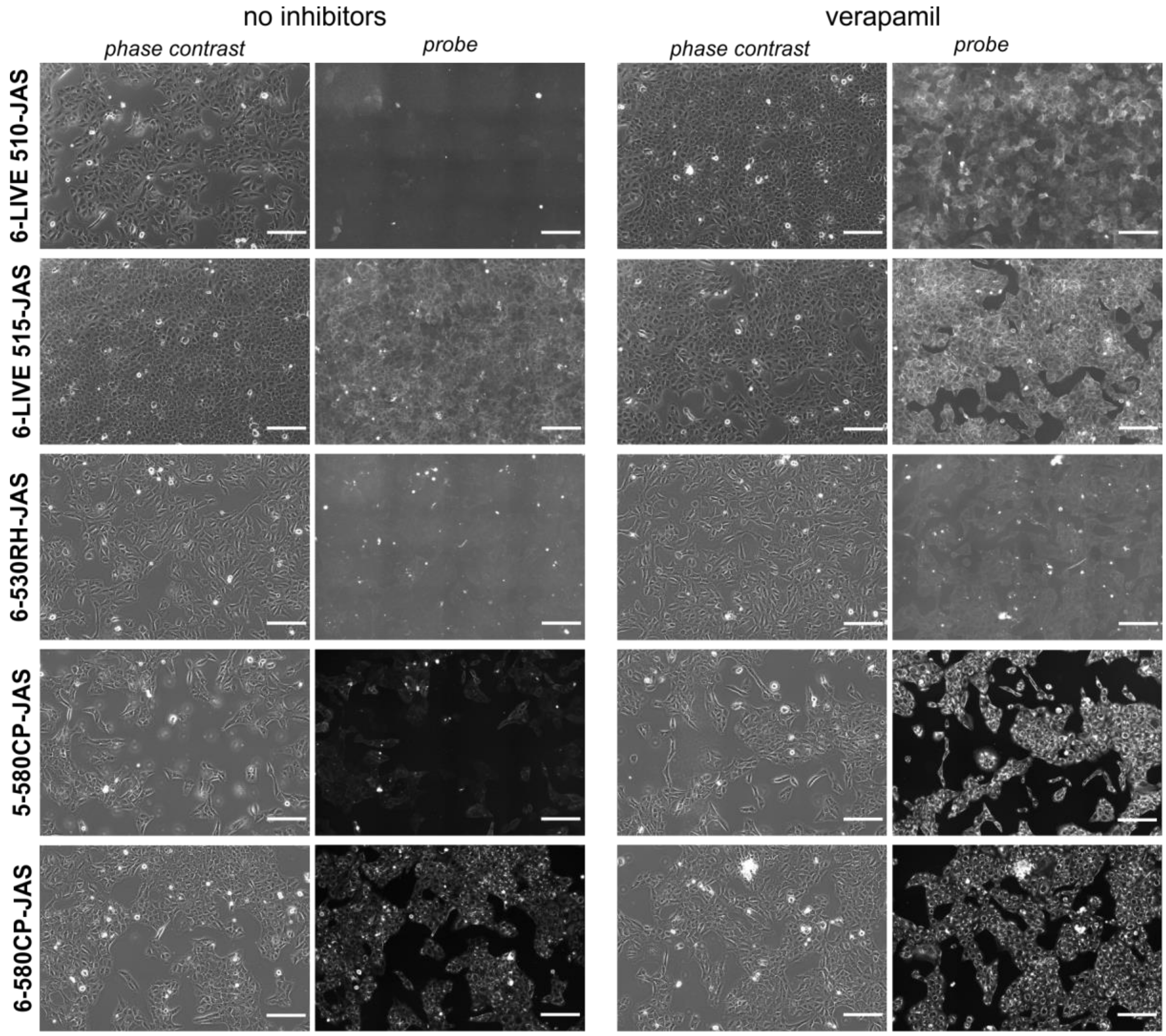

**Figure S3. Requirement of verapamil for efficient staining of U2OS cells.**
The cells were stained with 250 nM probe for 2 h at 37°C, washed twice and imaged on a wide-field automatic microscope at 20× magnification. 6-530RH-JAS only stained the cells at 1 µM. Scale bars 200 µm.

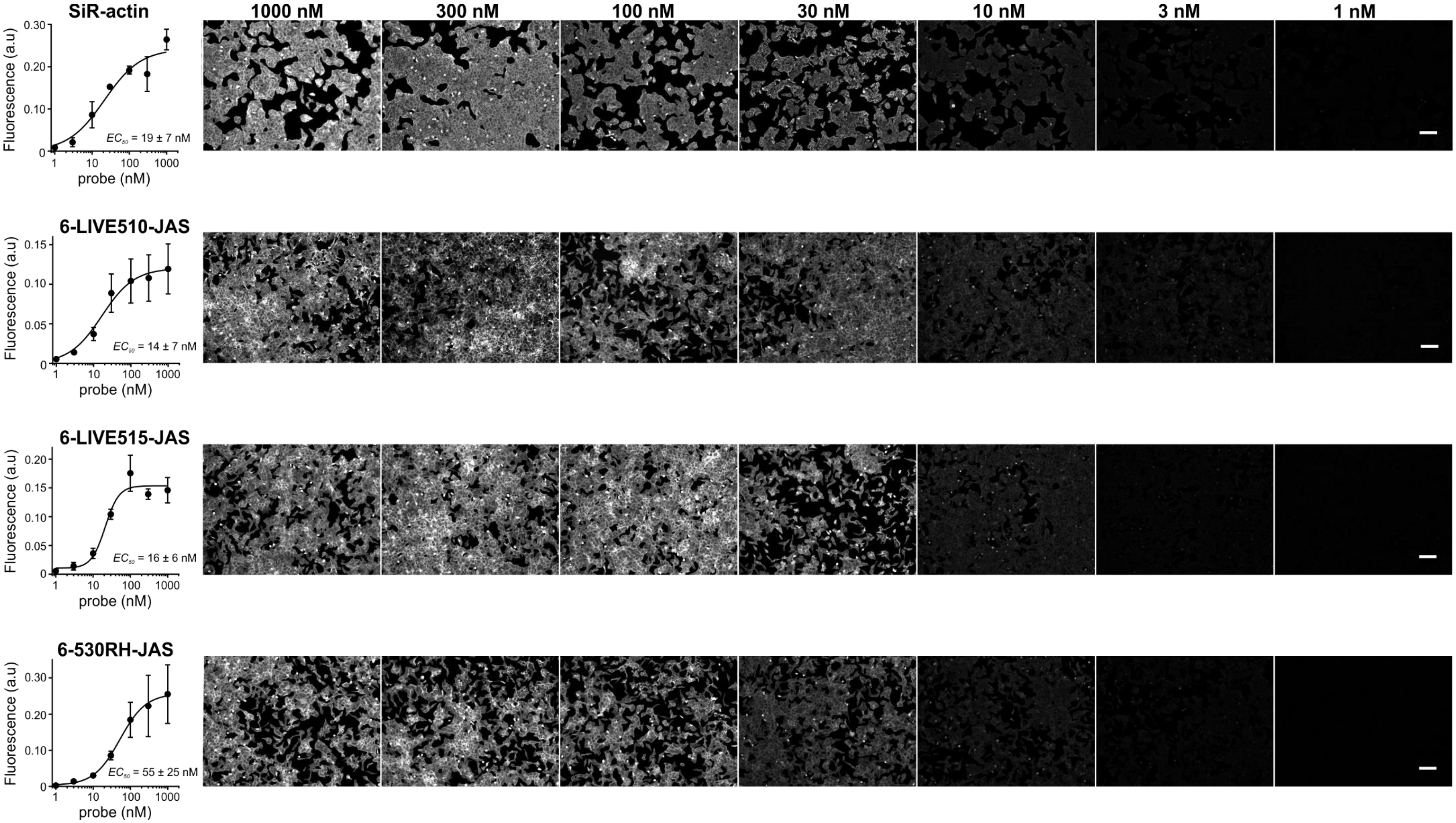

**Figure S4. Determination of $EC_{50}^{staining}$ (=$K_D^{F\text{-}actin}$) of probes by staining fixed and permeabilized U2OS cells**

The cells were fixed with formaldehyde, stained and imaged as described in the Methods section. Actin staining was quantified using an automated pipeline described in Figure S3. Background was defined as median pixel intensity in the areas not covered by cells. Background corrected images are shown. Scale bars 100 µm. The experiments were repeated 3 times, and the quantification of data is represented as mean ± SD. The fitting line is shown and the derived $EC_{50}$ is indicated.

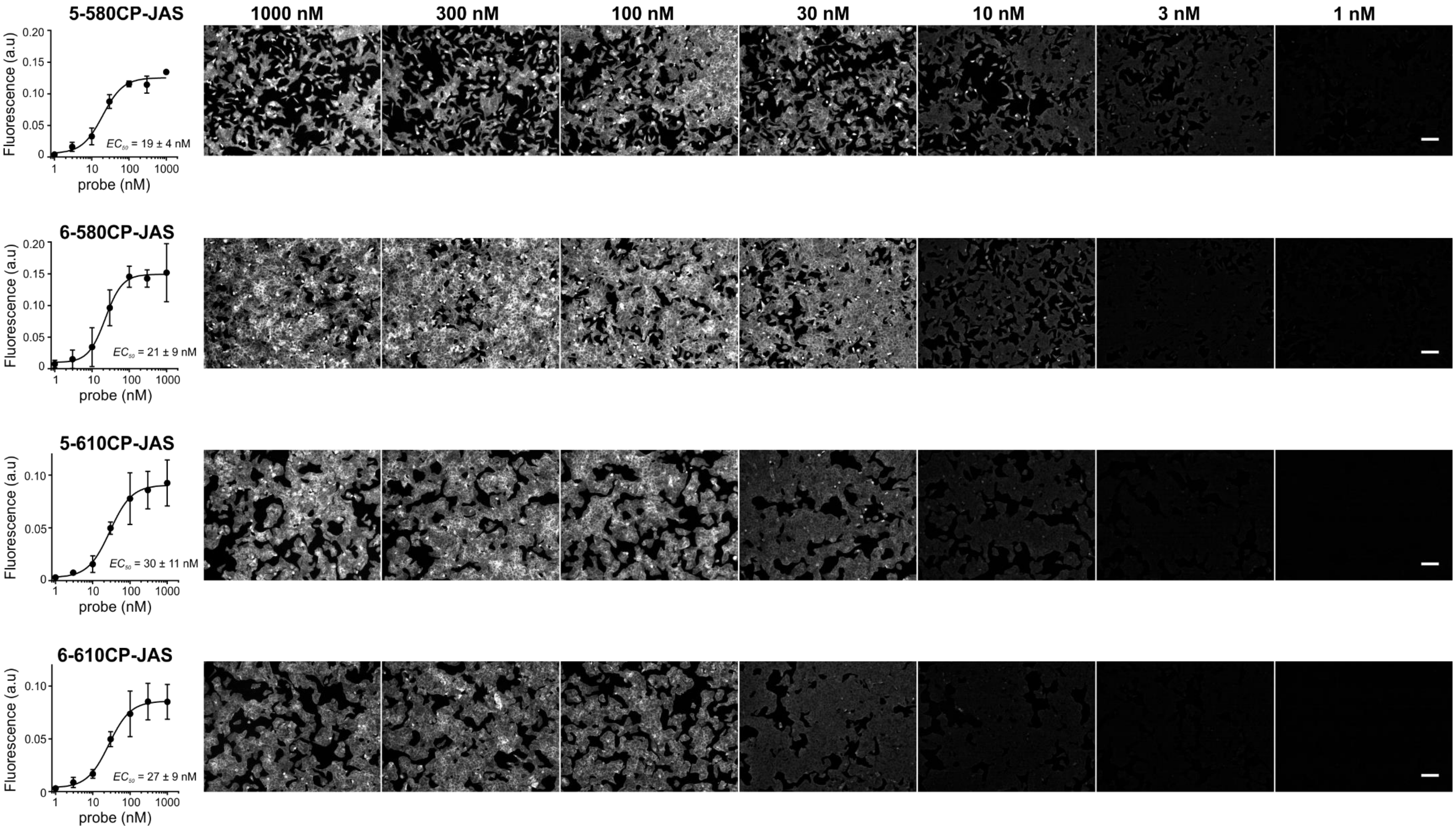

**Figure S4. Determination of EC50staining (=$K_D^{F\text{-}actin}$) of probes by staining fixed and permeabilized U2OS cells (continued).**

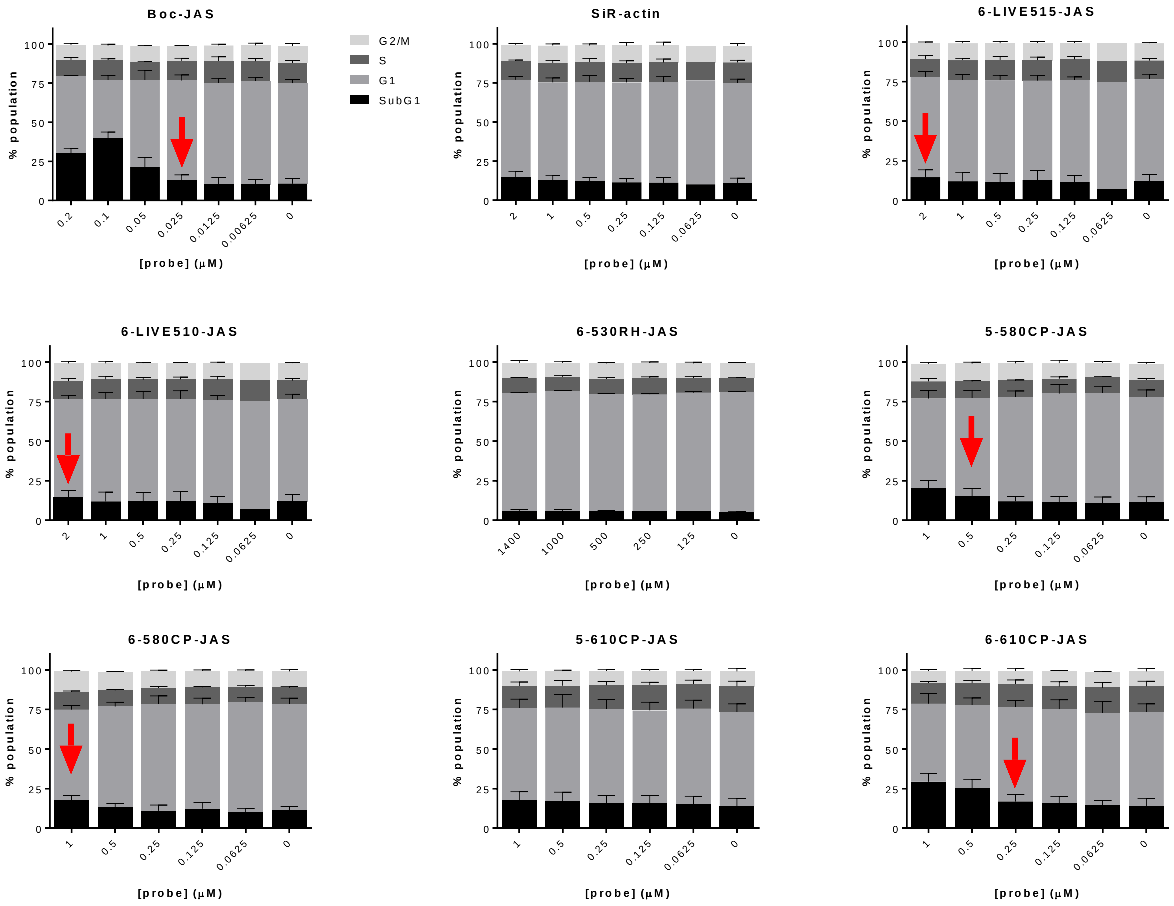

**Figure S5. Effect of the probes on the cell cycle progression analyzed by imaging cytometry.**
Hela cells were grown in the presence of the indicated probes for 24h and processed according to NucleoCounter® NC-3000™ (*Chemometec*) two-step cell cycle analysis protocol as previously described[1]. In each sample DNA content of ~10.000 cells in total were measured and the obtained cell cycle histograms were analysed with ChemoMetec NucleoView NC-3000 software, version 2.1.25.8. All experiments were repeated three times and the results are presented as means with standard deviations. Probe toxicity thereshold is defined as a minimum concentration at which increase in $SubG_1$ subpopulation can be detected. These values are marked with red arrows and listed in main text Table 1.

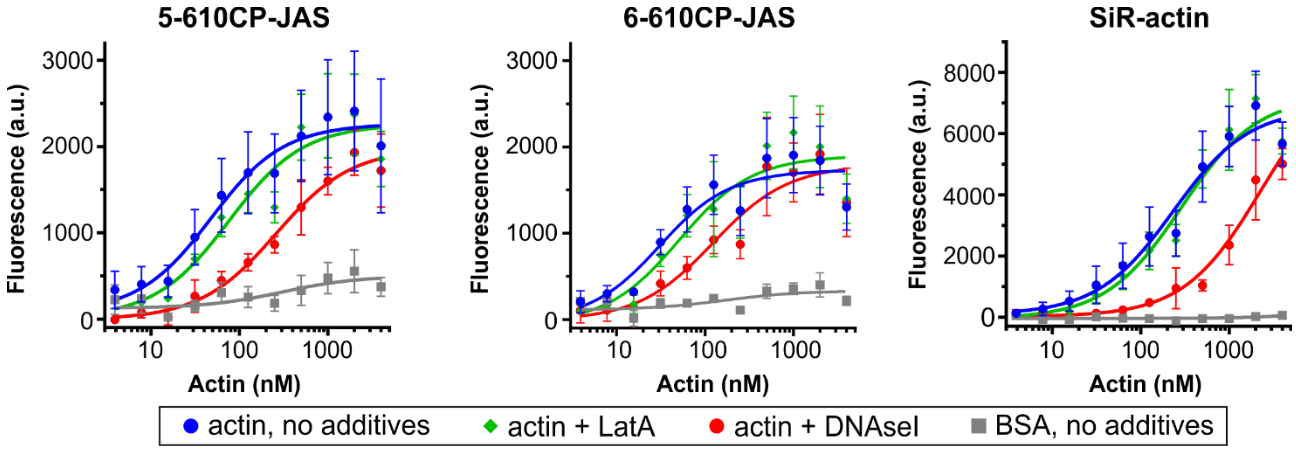

**Figure S6. Probe interaction with actin in a low salt buffer.**

Titration curves of 10 nM probes with increasing concentration of G-actin or BSA in the presence or absence of 5 µM latrunculin A (LatA) or 5 µM DNaseI. The data was fitted into a full equation of a single site binding. Data points are shown as mean ± SD; N = 5 without additives, N = 3 in the presence of actin polymerization inhibitors. The derived $K_D$ values are presented in main text Figure 3(a).

#### Supplementary Movies

**Video S1. Time-lapse Airyscan movie of a living fibroblast stained with 250 nM 6-610CP-JAS.**
Imaging was performed without washing after labeling for 2h at 37°C in DMEM containing 10% FBS. Scale bar 5 µm.

**Video S2. 3D STED rotating maximum intensity projections of a living fibroblast stained with 3 µM 6-610CP-JAS.**
Cells were stained for 1h at 37°C in DMEM containing 10% FBS and imaged after washing once with the same medium. Four projection represent raw and deconvolved (SVI Huygens) images acquired on microscope operating in confocal and STED modes. Scale bar 2 µm.

#### Supplementary Tables

**Table S1.** Optical properties of the probes.

| Compound | $\varepsilon^{dye}$ ($M^{-1}\cdot cm^{-1}$)[2-3] | $QY^{probe}_{SDS}$ |
|---|---|---|
| **6-LIVE510-JAS (10)** | 74000 | 0.756 ± 0.007 |
| **6-LIVE515-JAS (11)** | 56000 | 0.851 ± 0.008 |
| **6-530RH-JAS (12)** | 56000 | 0.849 ± 0.002 |
| **5-580CP-JAS (13)** | 90000 | 0.611 ± 0.004 |
| **6-580CP-JAS (14)** | 90000 | 0.624 ± 0.004 |
| **5-610CP-JAS (15)** | 100000 | 0.618 ± 0.006 |
| **6-610CP-JAS (16)** | 100000 | 0.615 ± 0.005 |
| **SiR-actin (17)** | 93000 | 0.509 ± 0.005 |

**Table S2.** Parameters of image acquisition on a wide-field Lionheart FX Automated Microscope

| Fluorophore | LED | Filter cube | Cells/objective | LED intensity | Integration time, ms | Camera gain |
|---|---|---|---|---|---|---|
| **Hoechst** | 365 nm | DAPI | living / 20× | 7 | 25 | 24 |
| | | 377/447 | fixed / 4× | 7 | 50 | 28 |
| **LIVE 510** | 465 nm | GFP | living / 20× | 7 | 75 | 30 |
| | | 469/525 | fixed / 4× | 8 | 75 | 28 |
| **LIVE 515/** | 523 nm | RFP | living / 20× | 10 | 500 | 30 |
| **530RH,** | | 531/593 | fixed / 4× | 8 | 200/75 | 28 |
| **580CP, 610CP** | 590 nm | Texas red | living / 20× | 7 | 100 | 28 |
| | | 586/647 | fixed / 4× | 8 | 100 | 30 |
| **SiR** | 623 nm | Cy5 | living / 20× | 7 | 50 | 24 |
| | | 628/685 | fixed / 4× | 8 | 150 | 30 |

**Table S3.** High resolution mass spectrometric characterization of the probes.

| Compound | Calculated m/z | Measured m/z | $\Delta_{calculated - measured}$ |
|---|---|---|---|
| **6-LIVE510-JAS (10)** | 1343.5422 | 1343.5412 | 0.0010 |
| **6-LIVE515-JAS (11)** | 1207.5674 | 1207.5680 | -0.0006 |
| **6-530RH-JAS (12)** | 1419.6134 | 1419.6072 | 0.0062 |
| **5-580CP-JAS (13)** | 1197.6383 | 1197.6377 | 0.0006 |
| **6-580CP-JAS (14)** | 1197.6383 | 1197.6384 | -0.0001 |
| **5-610CP-JAS (15)** | 1225.6696 | 1225.6705 | -0.0009 |
| **6-610CP-JAS (16)** | 1225.6696 | 1225.6694 | 0.0002 |
| **SiR-actin (17)** | 1241.6465 | 1241.6475 | -0.0010 |

### Supplementary materials and methods

#### Materials

Probes, verapamil (Sigma, Cat. No. V4629) and latrunculin A (Tocris, Cat. No. 3973) were dissolved in DMSO as 1000× stocks and were kept at -20°C. Bovine pancreas DNaseI was from Invitrogen (Cat. No. 18047019). The exact protein concentration was determined using Qubit™ 4 Protein Assay Kit (Thermo Scientific).

#### Determination of Quantum Yields

Relative quantum yields of the probes bound to actin were calculated by recording absorbance and fluorescence spectra using a multiwell plate reader Spark® 20M and glass bottom 96-well plates (MatTek, Cat. No. PBK96G-1.5-5-F) at room temperature (25 °C). To account for background due to light scattering, spectra of the solutions containing no probes, but equivalent amount of DMSO, were acquired and subtracted from the respective probe spectra. Absolute quantum yields of probes were measured using Quantaurus-QY absolute PL quantum yield spectrometer for samples containing 1 µM probe and 0.1 % SDS in PBS. Samples were incubated for 1 h at room temperature before measurement. The experiment was repeated three times in triplicates and the obtained $QY^{probe}{}_{SDS}$ are presented as mean ± SD in Supplementary Table S2. Relative quantum yields ($QY_{actin}$) of the probes bound to the target were calculated in a|e - UV-Vis-IR Spectral Software v2.2 (FluorTools) using the following formula (1):

$$QY_{actin} = QY_{SDS} \cdot \frac{A_{SDS} \cdot Fl_{actin}}{Fl_{SDS} \cdot A_{actin}}. \qquad (1)$$

where $QY_{SDS}$- absolute quantum yield of the probe dissolved in PBS containing 0.2 % SDS, $A_{SDS}$ – absorbance of the probe solution in PBS containing 0.1 % SDS at $\lambda_{max}$, $A_{actin}$ – absorbance of the probe in the solution containing excess of actin, $Fl_{SDS}$ – integrated fluorescence intensity of the probe solution in PBS containing 0.1 % SDS at $\lambda_{max}$, $Fl_{actin}$ – integrated fluorescence intensity of the probe in the solution containing excess of actin.

#### Preparation of G-actin

Lyophilized rabbit skeletal muscle actin (unlabeled and pyrene-labeled) was obtained from Cytoskeleton Inc (Cat. No. AKL99). For preparing G-actin, 1 mg of actin was dissolved in 0.5 ml General Actin Buffer (5 mM Tris·HCl, 0.2 mM $CaCl_2$), supplemented with 0.2 mM ATP, and incubated on ice for 1 h, in order to depolymerize the oligomers that might have formed during storage. Any remaining oligomers or aggregates were removed by centrifugation at 14000 rpm for 30 min. at 4°C. The solution was transferred into a new tube and kept on ice. This gave 46.5 µM actin stock solution.

#### Polymerization of pyrene-actin

4 µM pyrene-actin was prepared in General Actin Buffer, supplemented with 0.2 mM ATP and 0.5 mM DTT containing 10 nM SiR-actin or 610CP-JAS probes or 0.1% DMSO. Aliquots of 50 µl were added to a black 96-well half-area plate and the fluorescence was read for 3 min. at 37°C, taking one read every 30 s. Where required, 5 µl of 10× actin polymerization buffer was quickly added and the measurements were continued for 60 min. Pyrene fluorescence was measured with Tecan Spark20M plate reader in a top-reading mode, using excitation of 360 nm (bandwidth 20 nm) and recording emission at 410 nm (bandwidth 10 nm).

#### General experimental information for organic synthesis

All standard chemicals and solvents were purchased from commercial suppliers (Sigma-Aldrich, Merck, VWR) and used without further purification. Boc-jasplakinolide was synthesized via solid-phase-synthesis according to the previously described procedure by Milroy et al. [4].

Analysis was performed on a ThermoScientific Ultimate 3000 system with a diode arrey detector using a Kinetex 2,6µm C18 (4,6 x 75 mm) column at 20 °C. Eluent A: MeCN; Eluent B: $H_2O$ +0,05% v/v TFA. Purification was performed on an Interchim Puriflash 4250 system with a TOYDAD800 four channel UV-Vis-detector using an Interchim puriFlash 5µm C18 AQ (21.2 x 250 mm) column. Eluent A: MeCN; Eluent B: H2O +0,1% v/v TFA

Low resolution mass spectra (50 – 3500 m/z) was performed with electrospray ionization (ESI-MS) technique using a Varian 500-MS spectrometer. High resolution mass spectra (ESI-HRMS) of the jasplakinolide conjugates were obtained on a Bruker micro TOF (ESI-TOF-MS) spectrometer.

#### Synthesis

*Boc-deprotection of Boc-Jasplakinolide*:

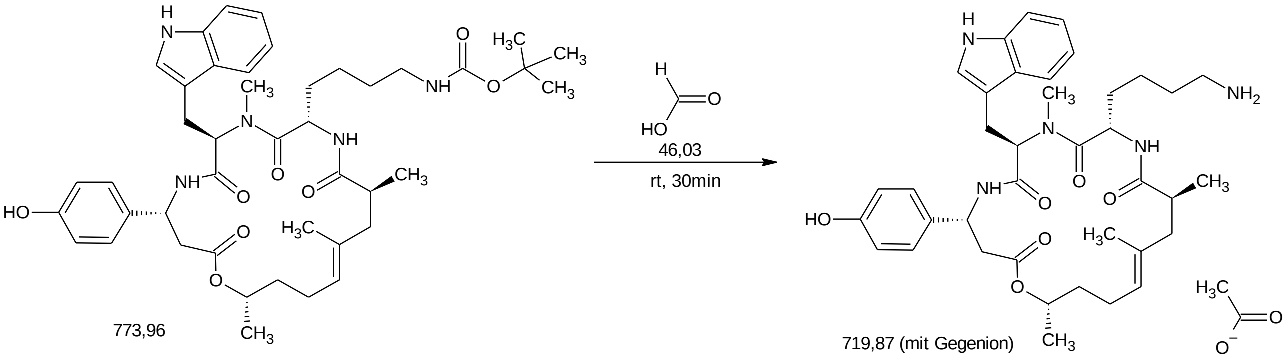

2.0 mg (2.58 µmol) Boc-Jasplakinolide was dissolved in 210 µl Formic acid and stirred at room temperature for 30 minutes. The reaction was controlled via analytical HPLC. After completion of the reaction, 17 µl ultrapure water were added, the reaction mixture frozen on dry ice and lyophilized. To remove all traces of formic acid, another 170 µl ultrapure water were added, the reaction mixture frozen and lyophilized again, yielding 1.86 mg (quant.) of jasplakinolide amine formiate salt.

*Linker attachement*:

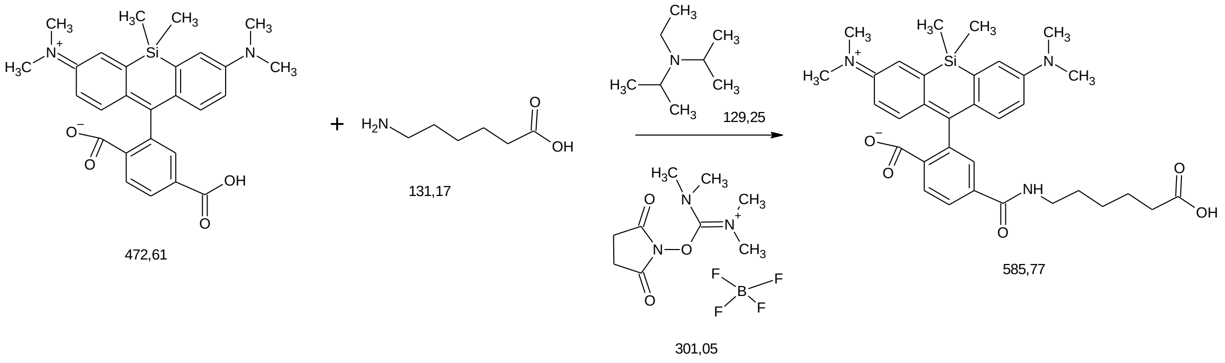

2.0 mg of dye-COOH were dissolved under argon atmosphere in an oven dried pear shaped flask in 400 µl abs. DMSO. 26 eq of N,N-Diisopropylethylamine and 1.2 eq of TSTU were added and the reaction mixture was stirred for 5 minutes under inert gas. After in situ formation of the NHS-ester, 3.8 eq of 6-aminohexanoic acid were added, the reaction mixture was sonicated for 15 minutes. After addition of 20 µl ultrapure water, stirring at room temperature was continued for 15 minutes and the conversion monitored by analytical HPLC. The reaction mixture was quenched, adding 30 eq of formic acid, frozen on dry ice and lyophilized, followed by preparative RP-HPLC, yielding dye-C6-linker in 75-85% yield.

*NHS-ester formation*:

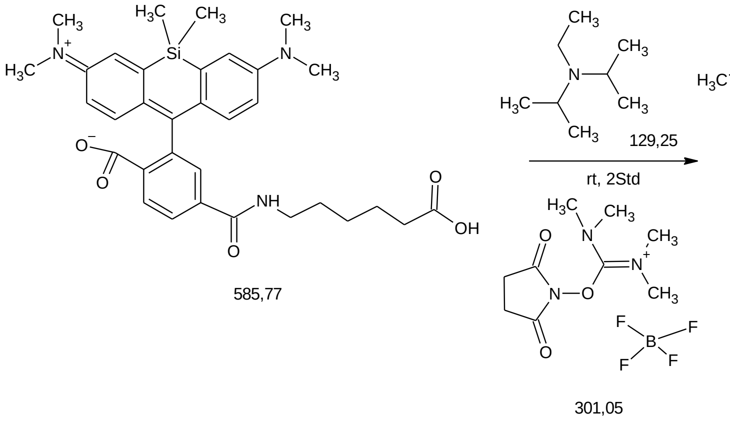

1.5 mg of dye-C6-linker were dissolved under argon atmosphere in an oven dried pear shaped flask in 300 µl of abs. DMSO. 8 eq of N,N-Diisopropylethylamine and 2.6 eq of TSTU were added and the reaction mixture was stirred for 60 minutes under inert gas. The reaction conversion was monitored by analytical HPLC. After full conversion, the reaction mixture was frozen on dry ice, lyophilized and purified via preparate HPLC, yielding dye-C6-linker-NHS in 60-80% yield.

*Jasplakinolide coupling*:

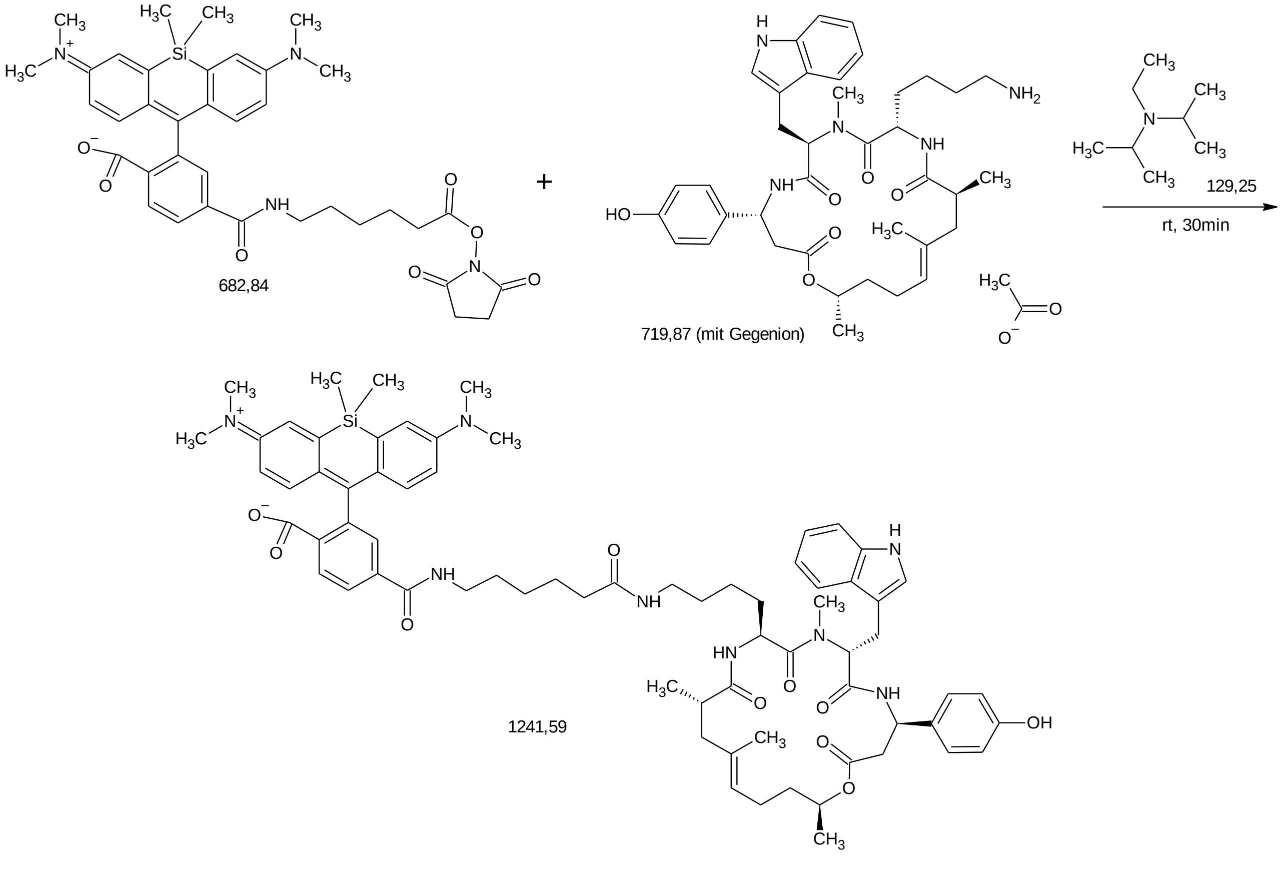

1 mg of dye-C6-linker-NHS were dissolved under argon atmosphere in an oven dried pear shaped flask in 145 µl abs. DMSO. 8 eq of N,N-Diisopropylethylamine and 1 – 1.2 eq of jasplakinolide-amine formiate salt were added and the reaction mixture was stirred for 40 minutes under inert gas. The reaction conversion was monitored by analytical HPLC.After full conversion, the reaction mixture was frozen on dry ice, lyophilized and purified via preparative HPLC, yielding dye-C6-linker-jasplakinolide in 40-70% yield.

#### HPLC traces, UV absorbance and HRMS-ESI spectra

HPLC trace of 6-LIVE 510-JAS (**10**)

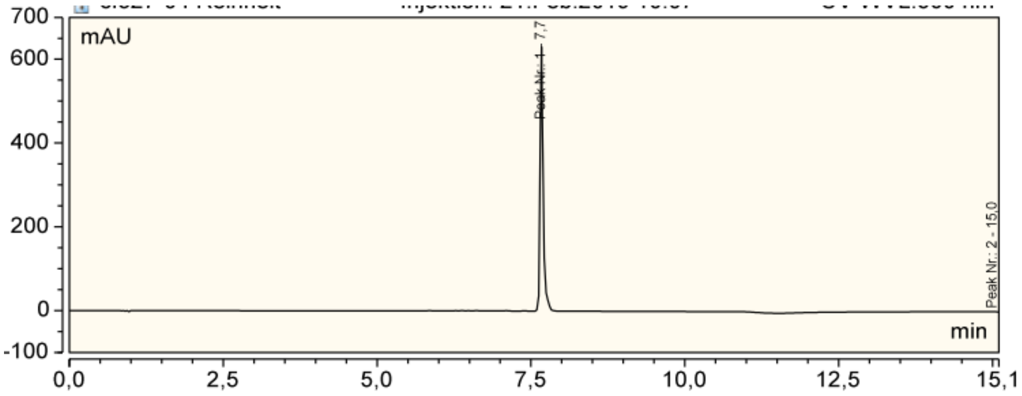

Absorbance spectrum at HPLC peak of 6-LIVE 510-JAS (**10**).

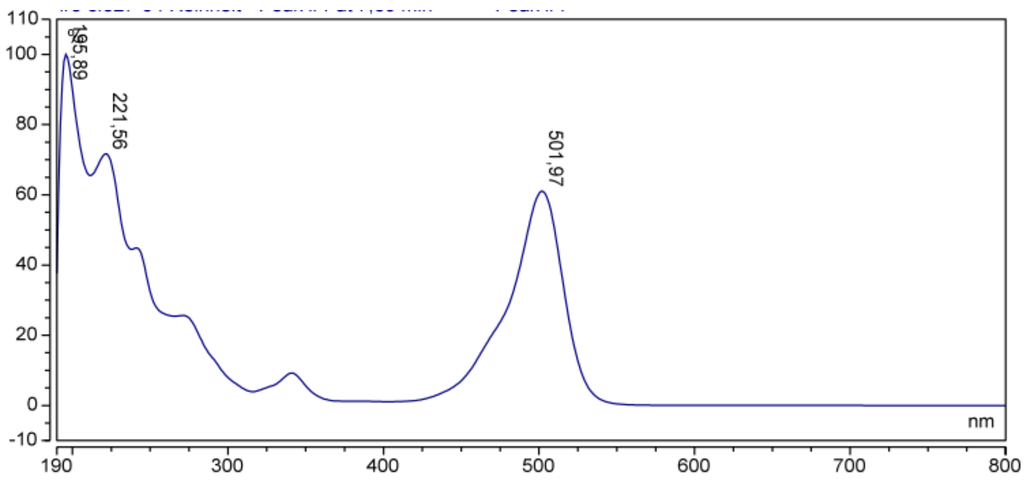

Measured and calculated HRMS-ESI spectra of 6-LIVE 510-JAS (**10**)

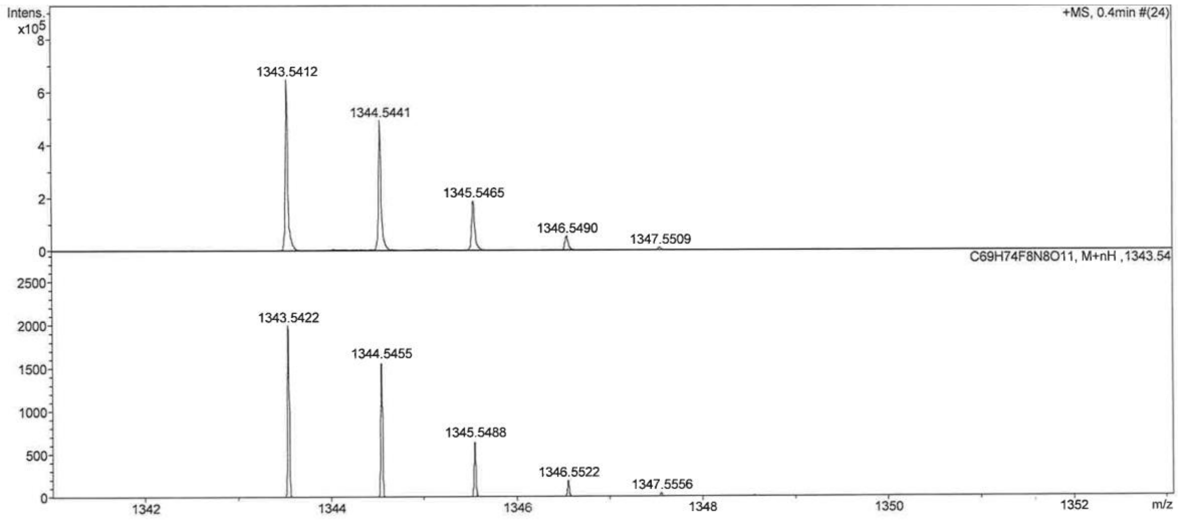

HPLC trace of 6-LIVE 515-JAS (**11**)

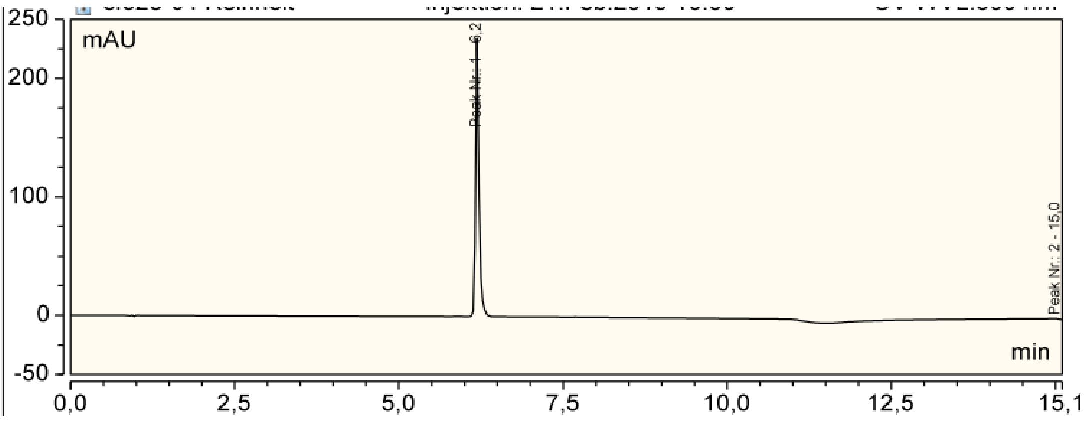

Absorbance spectrum at HPLC peak of 6-LIVE 515-JAS (**11**).

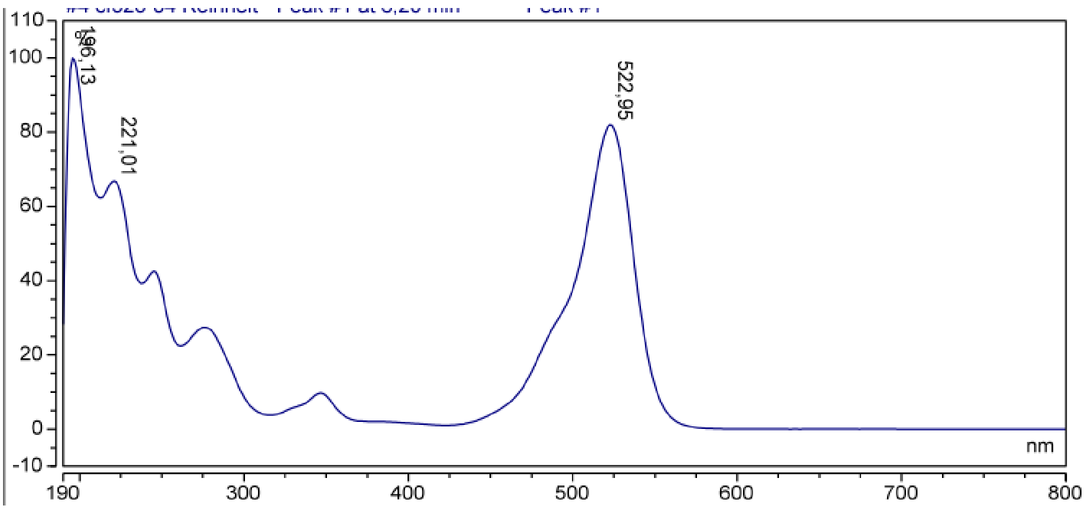

Measured and calculated HRMS-ESI spectra of 6-LIVE 515-JAS (**11**)

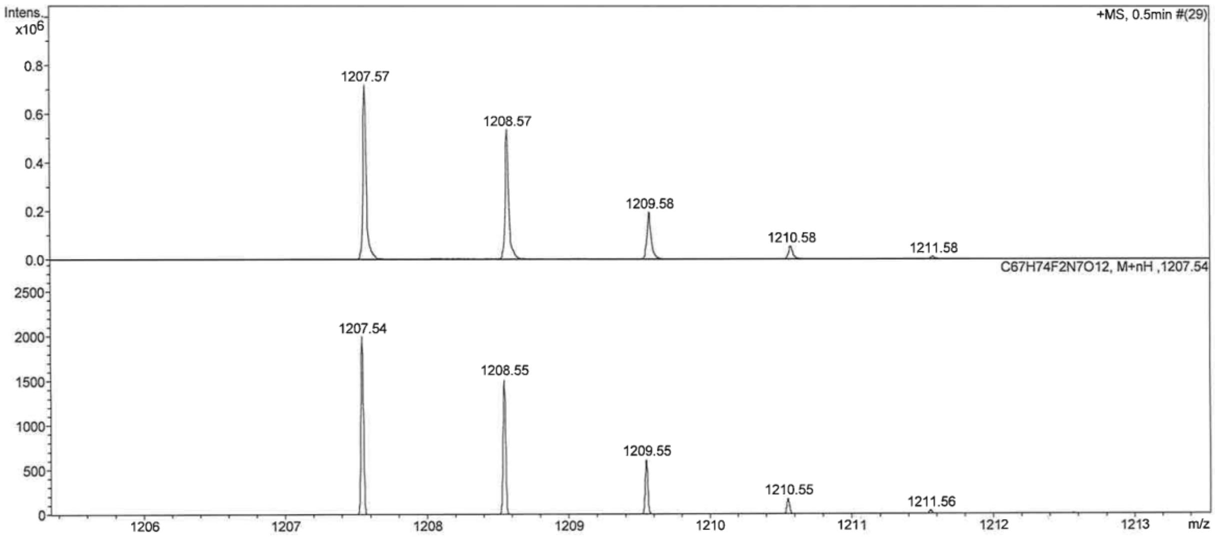

HPLC trace of 6-530RH-JAS (**12**).

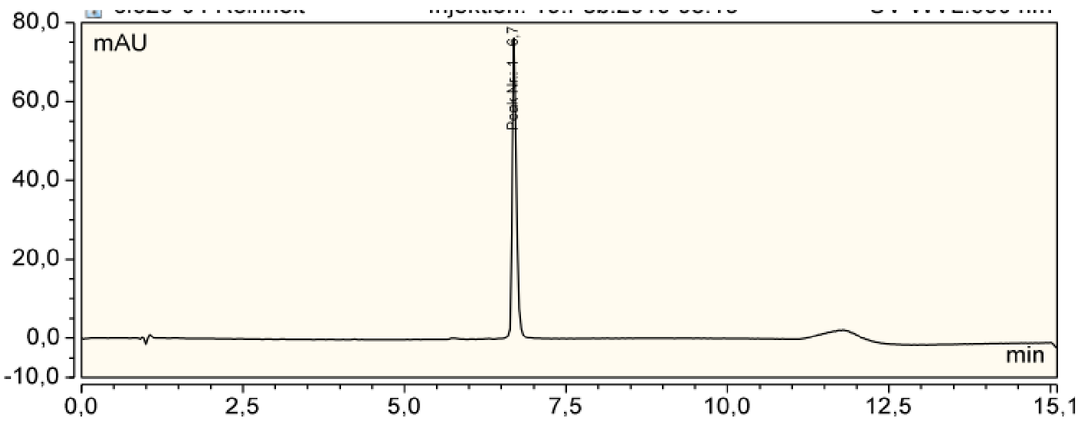

Absorbance spectrum at HPLC peak of 6-530RH-JAS (**12**).

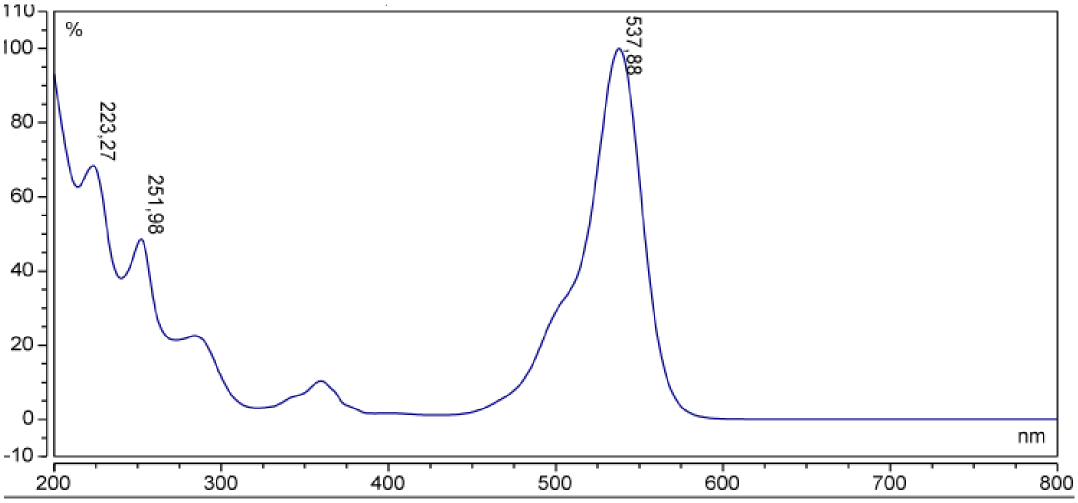

Measured and calculated HRMS-ESI spectra of 6-530RH-JAS (**12**).

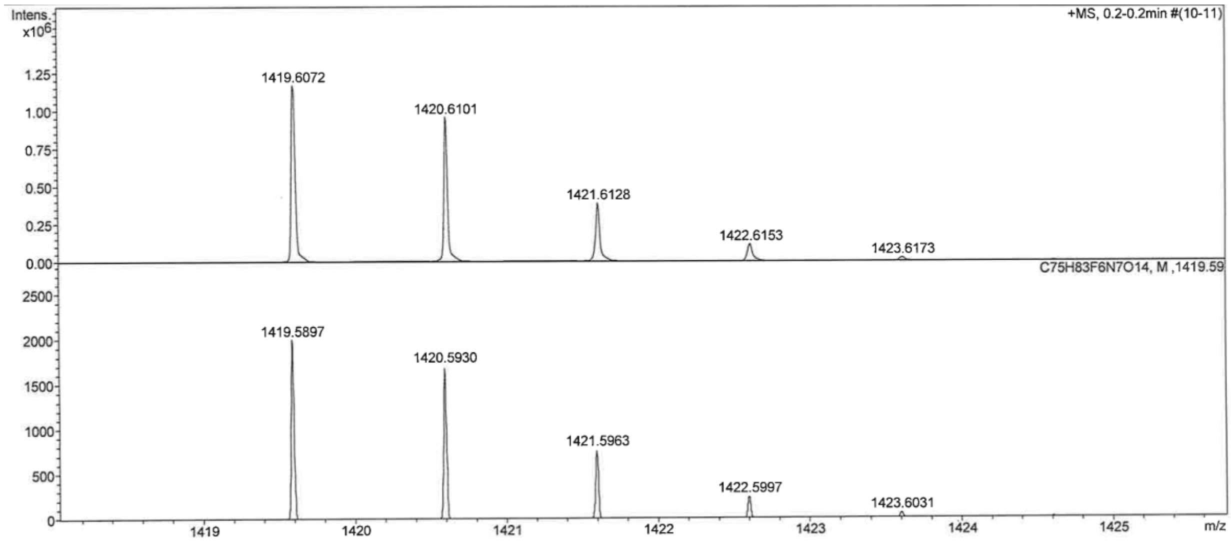

HPLC trace of 5-580CP-JAS (**13**).

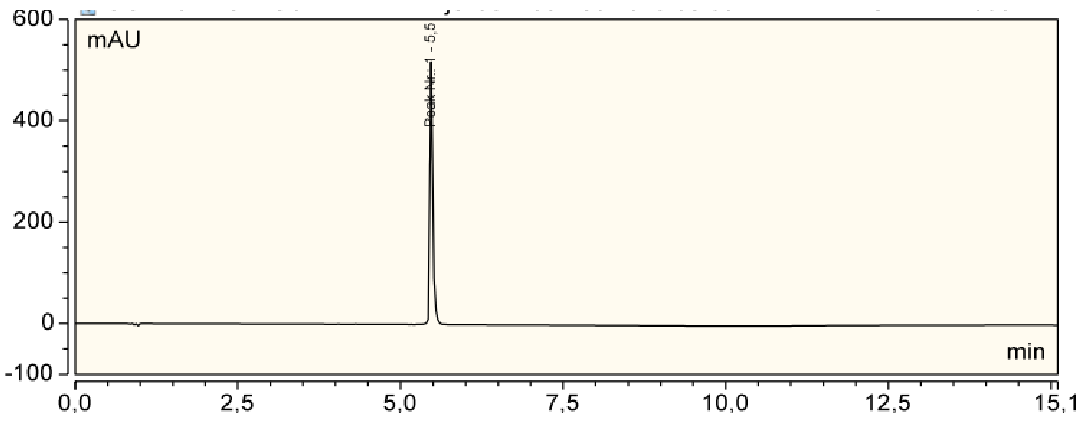

Absorbance spectrum at HPLC peak of 6-580CP-JAS (**13**).

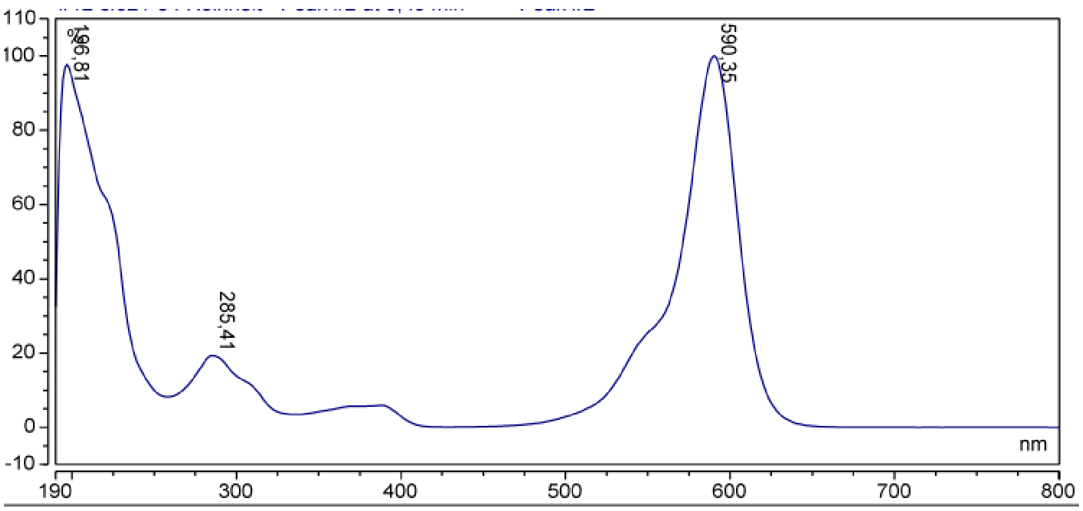

Measured and calculated HRMS-ESI spectra of 5-580CP-JAS (**13**).

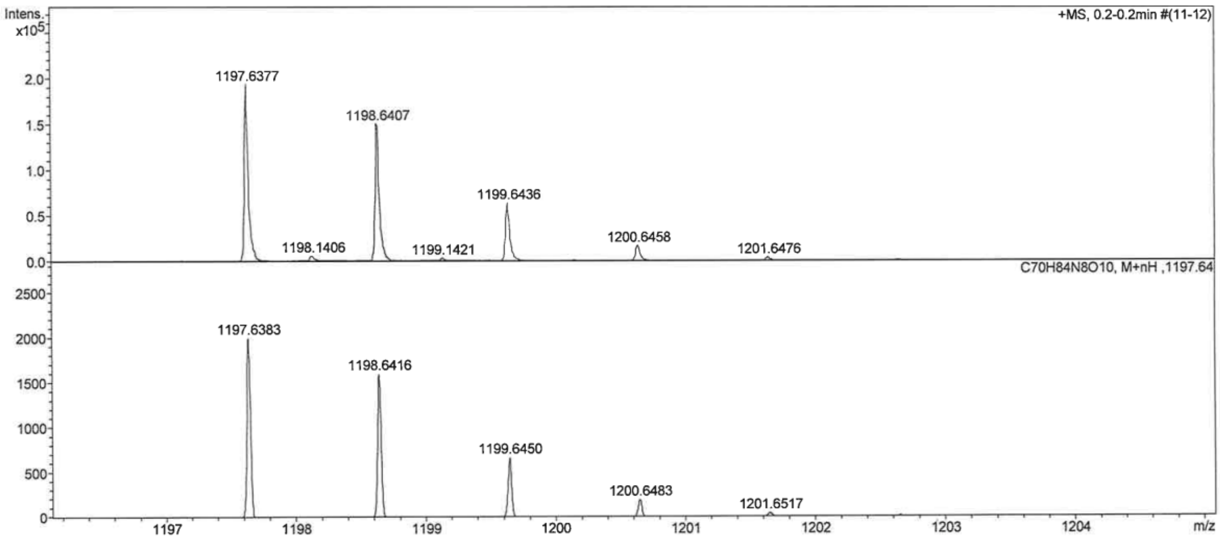

HPLC trace of 6-580CP-JAS (**14**).

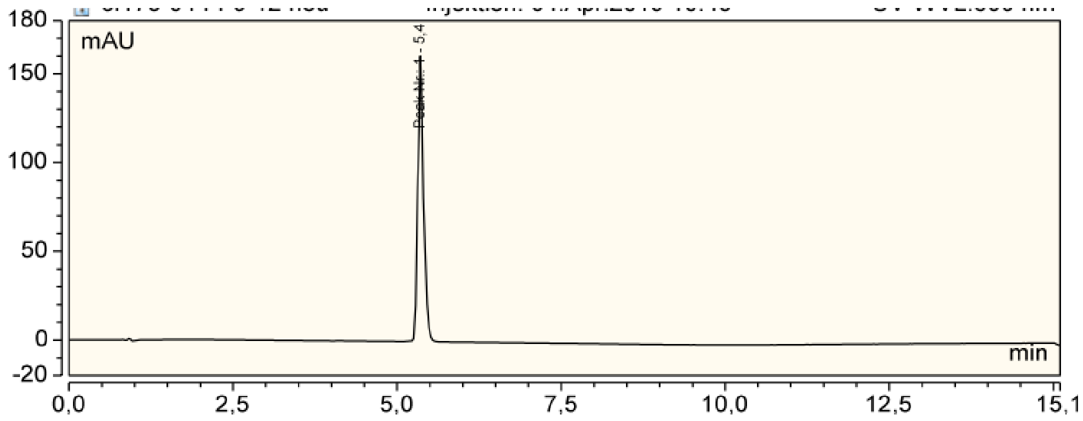

Absorbance spectrum at HPLC peak of 6-580CP-JAS (**14**).

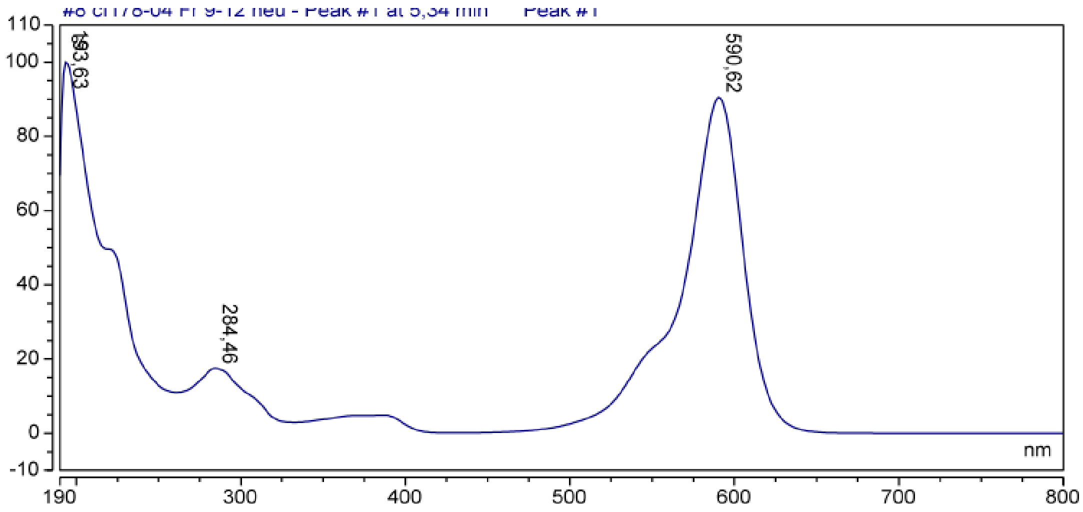

Measured and calculated HRMS-ESI spectra of 6-580CP-JAS (**14**).

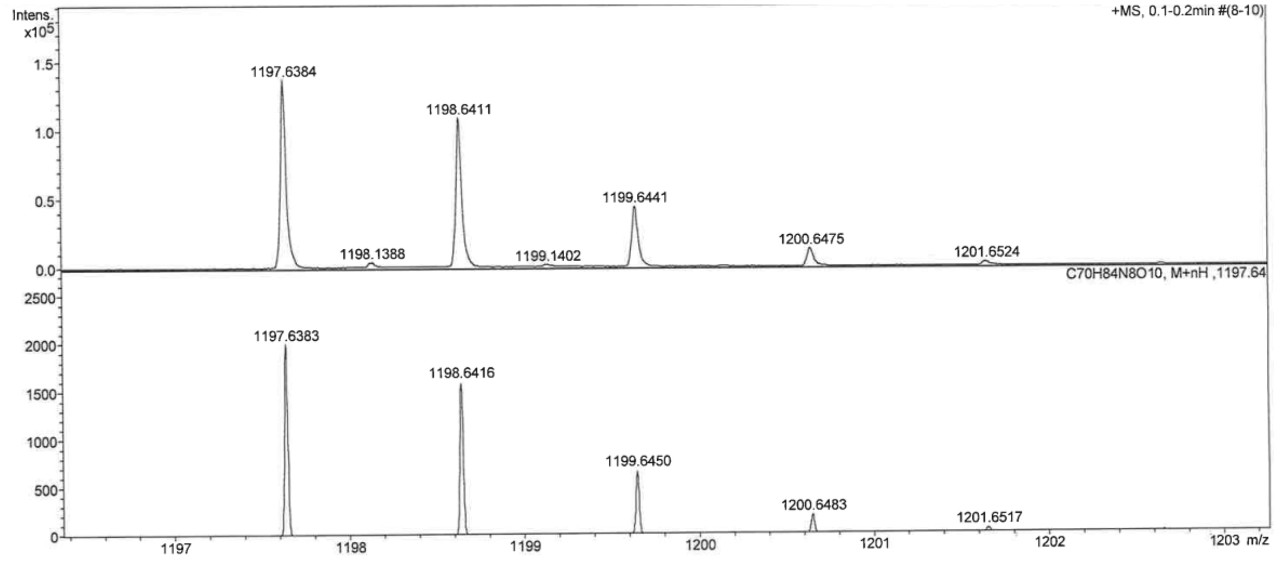

HPLC trace of 5-610CP-JAS (**15**).

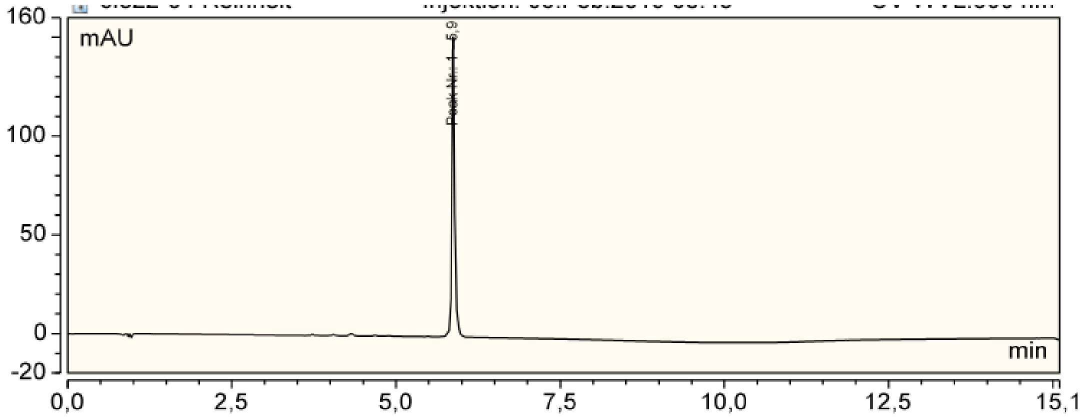

Absorbance spectrum at HPLC peak of 5-610CP-JAS (**15**).

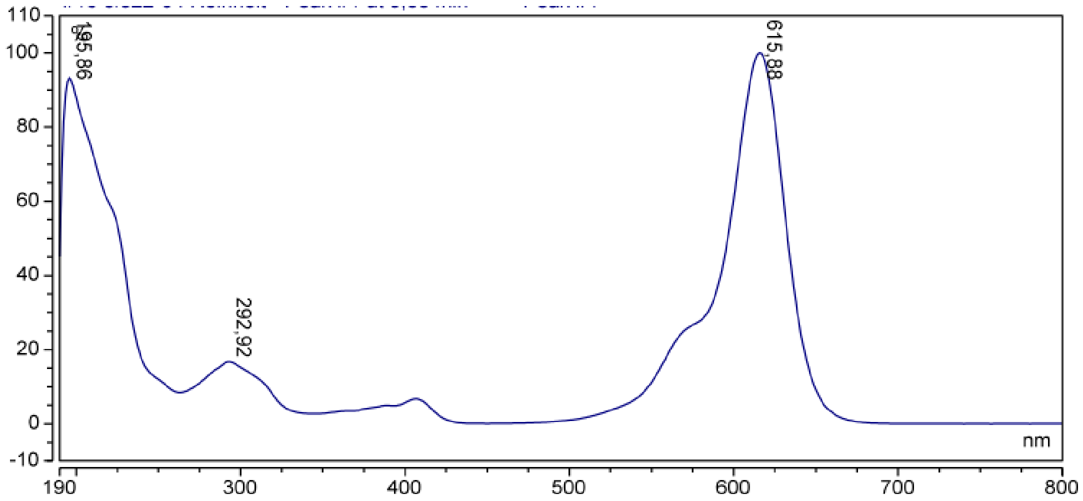

Measured and calculated HRMS-ESI spectra of 5-610CP-JAS (**15**).

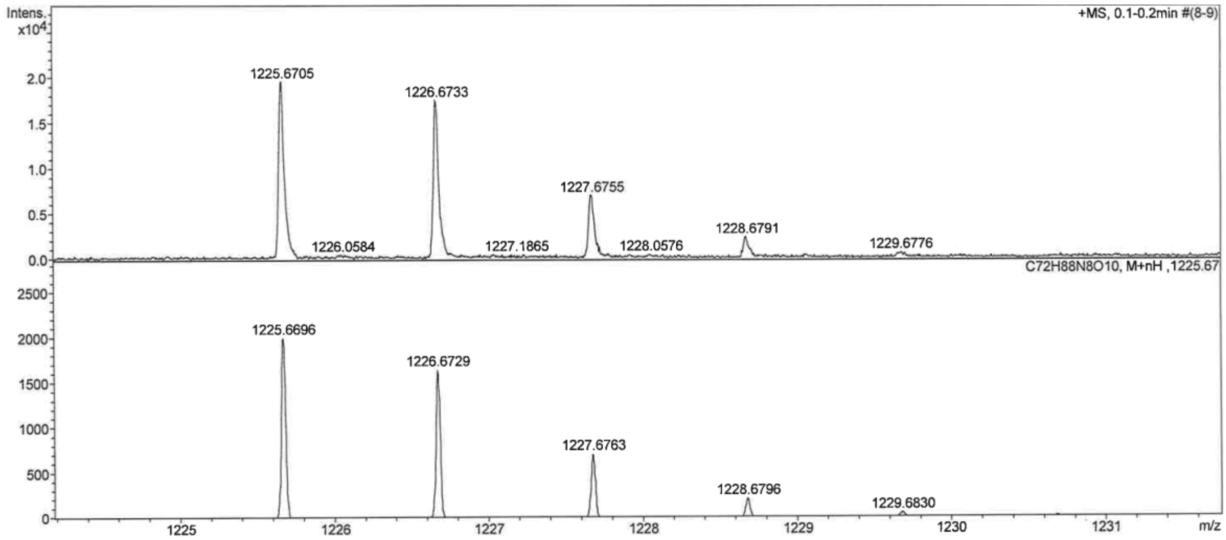

HPLC trace of 6-610CP-JAS (**16**).

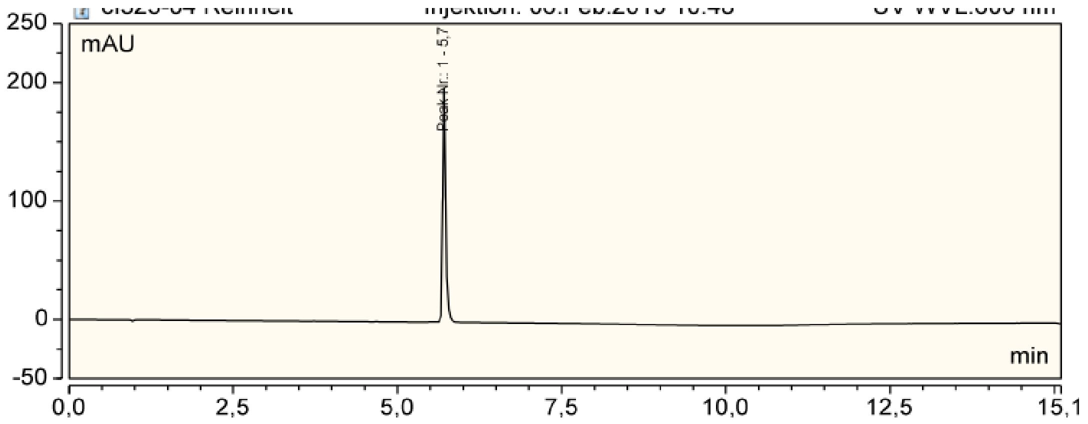

Absorbance spectrum at HPLC peak of 6-610CP-JAS (**16**).

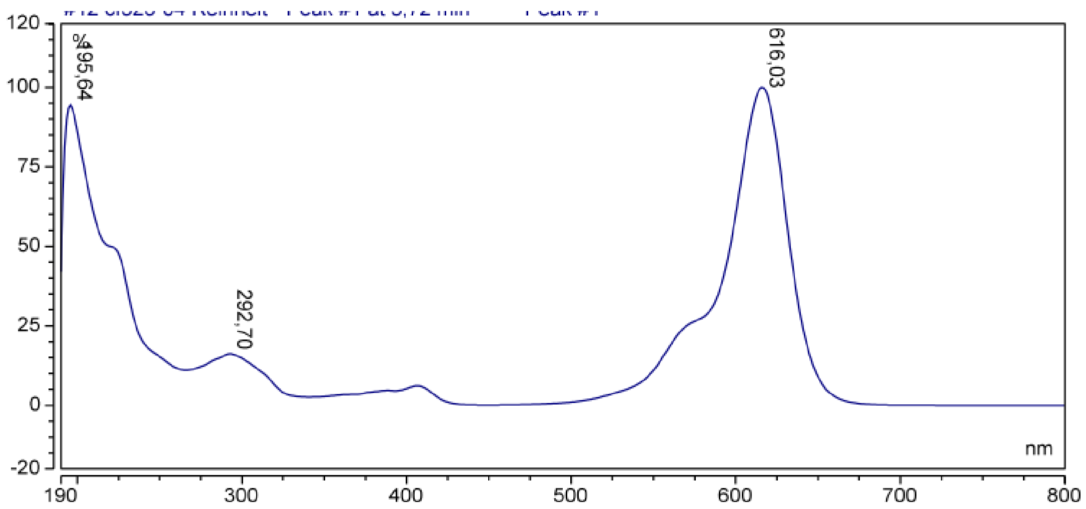

Measured and calculated HRMS-ESI spectra of 6-610CP-JAS (**16**).

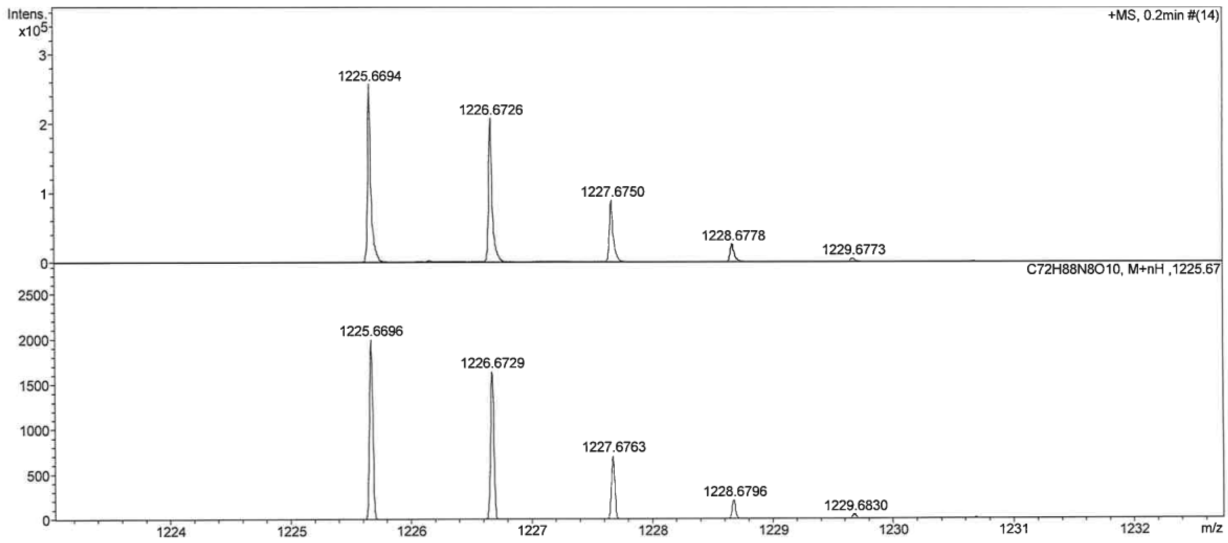

HPLC trace of SiR-actin (**17**).

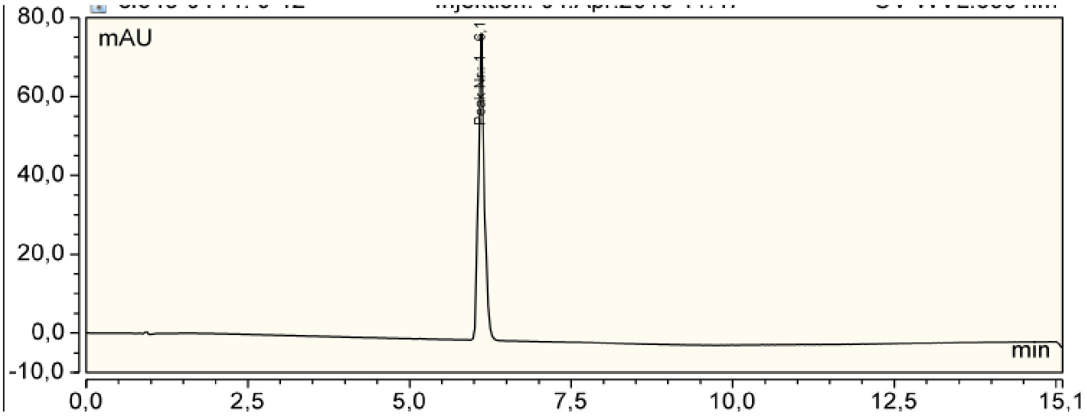

Absorbance spectrum at HPLC peak of SiR-actin (**17**).

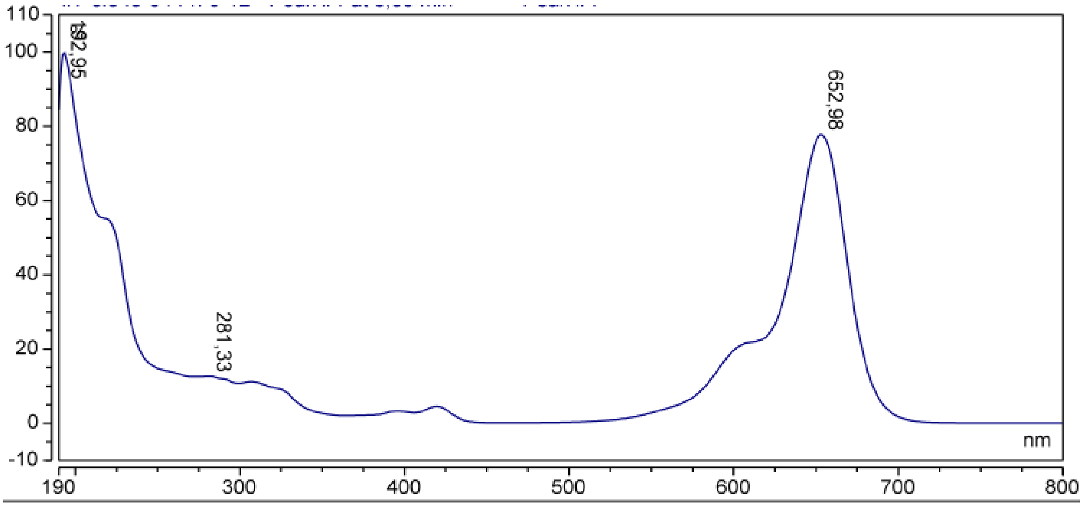

Measured and calculated HRMS-ESI spectra of SiR-actin (**17**).

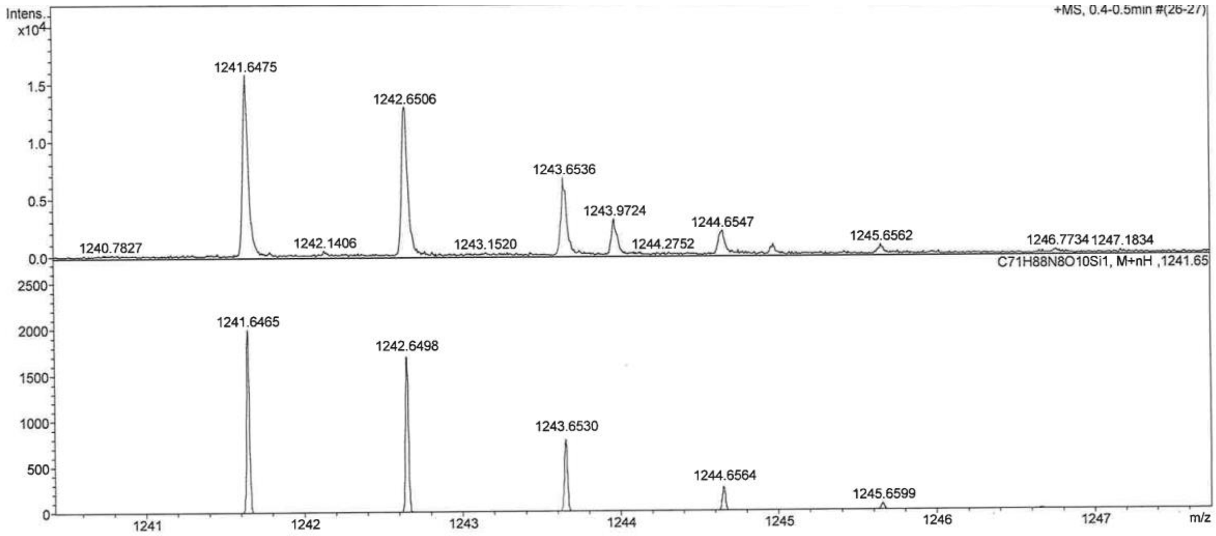
